# Reactive persistence of riverine metapopulations

**DOI:** 10.64898/2025.12.02.691852

**Authors:** Lorenzo Mari, Enrico Bertuzzo, Andrea Rinaldo, Marino Gatto, Renato Casagrandi

## Abstract

Understanding the conditions that favor the persistence of metapopulations inhabiting riverine landscapes (riverscapes) is critical to guiding conservation and restoration efforts aimed at preserving biodiversity and ecological integrity in freshwater environments. In this study, we present a modeling framework to examine the transient persistence of fluvial metapopulations—the temporary occupation of riverscape patches by a metapopulation expected to become extinct in the long run. Our approach is theoretically grounded in the concept of ecological reactivity, which effectively complements asymptotic stability analysis in the study of the short-term response of ecological systems to impulsive perturbations of otherwise stable steady states. Our results suggest that, under ecohydrological conditions conducive to reactive metapopulation extinction equilibria, a metapopulation asymptotically bound to extinction can still colonize parts of the riverscape for significant periods of time. We also find that, in the presence of repeated positive perturbations of the extinction equilibrium, the temporal scales associated with these transient phenomena may allow for reactive pseudo-persistence. This phenomenon involves an arbitrarily long delay in the eventual extinction of the metapopulation, which may occur well below the deterministic extinction threshold. Identifying the ecohydrological drivers of reactivity-driven metapopulation persistence and the riverscape patches that contribute most to transient metapopulation dynamics over different temporal scales may provide valuable suggestions for the spatial prioritization of conservation and restoration efforts.

## Introduction

River ecosystems constitute rich reservoirs of biodiversity, supporting a wide variety of species that have evolved to thrive in their dynamic and interconnected habitats (Fagan, 2002; Dudgeon et al., 2006; Rinaldo et al., 2020). These ecosystems host a broad range of organisms, from microscopic phytoplankton and aquatic invertebrates to fish and amphibians. The riparian zones along the river banks also contribute to this diversity by providing essential habitats for various terrestrial species, such as insects, reptiles, birds, and mammals (Sabo et al., 2005; Riis et al., 2020). This teeming biological diversity is related to vital ecosystem services, including nutrient cycling, pollution control, drinking water supply, and recreation (Petsch et al., 2023). From a landscape ecology perspective, river ecosystems are often fragmented, with various isolated habitats separated by natural or artificial barriers, but still connected by hydrological corridors (Fuller et al., 2015; Belletti et al., 2020). Thus, a metapopulation perspective (Casagrandi and Gatto, 1999; Hanski, 1999; Hanski and Ovaskainen, 2000; Rybicki and Hanski, 2013) should be applied to obtain an effective characterization of the ecological dynamics of fluvial species (Talluto et al., 2024). Metapopulations exhibit a dynamic interplay of local extinction and (re)colonization processes, where sub-populations may go extinct in one patch but be rescued via immigration from occupied ones (Levins, 1969). Specifically, riverine metapopulations (Mari et al., 2014) are driven by the dynamics of river systems, with their flow patterns, connectivity, and seasonal variations, to maintain genetic diversity and facilitate the colonization of empty patches (Rinaldo et al., 2020).

Despite their ecological importance, freshwater ecosystems are among the most vulnerable to the accelerating impacts of global change and anthropogenic pressures (Reid et al., 2019; Haase et al., 2023). Therefore, understanding the factors that affect the persistence of metapopulations that live in (or along) rivers, such as habitat connectivity, patch quality, and landscape structure, represents a critical challenge in guiding conservation and restoration efforts aimed at preserving biodiversity and ecological integrity (Altermatt, 2013). To date, much of the research on this topic has focused on long-term dynamics or even asymptotic persistence (e.g., Speirs and Gurney, 2001; Fagan, 2002; Pachepsky et al., 2005; Lutscher et al., 2010; Mari et al., 2014; Yeakel et al., 2014; Van Looy and Piffady, 2017; Terui et al., 2018; Ma et al., 2020; Bertassello et al., 2021; Giezendanner et al., 2021; Bertassello et al., 2022a,b; Nguyen et al., 2026). However, the continuous disturbance to which real metapopulations are subjected can make transient dynamics at least as important as asymptotic behavior (Hastings, 2004; Aiken et al., 2023), a concept that can be traced back to *r/K* selection theory (MacArthur and Wilson, 1967; Roughgarden, 1979; Matessi and Gatto, 1984). This remark is especially relevant to metapopulations that, although destined to disappear in the long run, could exhibit lengthy relaxation periods, thus creating an extinction debt (Tilman et al., 1994; Hanski and Ovaskainen, 2002; Kuussaari et al., 2009), or undergo a temporary but pronounced expansion, resulting in boom-and-bust dynamics (Simberloff and Gibbons, 2004; Strayer et al., 2017). In either case, the study of transient behavior may provide a more appropriate framework for evaluating ecological dynamics than long-term equilibrium analysis.

Stemming from Neubert and Caswell’s (1997) seminal paper, reactivity analysis and its generalizations (e.g., Verdy and Caswell, 2008; Mari et al., 2017) have emerged as an effective methodological framework for studying the short-term response of ecological systems to impulsive perturbations to otherwise asymptotically stable attractors (typically, equilibria). Formally, reactivity is defined as the maximum instantaneous rate at which perturbations to a stable steady state can be amplified; thus, an asymptotically stable equilibrium is reactive if there exist perturbations that can temporarily grow before eventually vanishing. Assessing the possible growth of perturbations requires the definition of a norm, a function that maps state-space coordinates into a non-negative scalar that can be interpreted as an appropriate measure of the distance from the reference steady state. Although the original formulation of the theory (Neubert and Caswell, 1997) relied on the Euclidean distance of the state vector from equilibrium to measure the amplification of perturbations, this metric (*ℓ*^2^-norm) lacks a clear ecological interpretation (Harrington et al., 2022). In contrast, measuring perturbations by their absolute deviation from the reference equilibrium (*ℓ*^1^-norm) may have a more straightforward biological interpretation, especially for positive perturbations of an extinction (null) equilibrium, whose *ℓ*^1^-norm simply corresponds to the sum of state variables (or a generic linear transformation thereof, see Mari et al., 2017). For this reason, the *ℓ*^1^-norm has been occasionally but effectively used to characterize transient dynamics in single-species ecological problems (Townley and Hodgson, 2008; Stott et al., 2010, 2011; Huang and Lewis, 2015; Harrington et al., 2022) and, more recently, in disease ecology and epidemiological applications (Trevisin et al., 2024; Mari et al., 2025; Trevisin et al., 2025).

In this study, we investigate the transient persistence of metapopulations inhabiting fluvial landscapes. To ensure our findings are robust to diverse topological structures, we use synthetic river networks that, while not tied to a specific case study, can realistically replicate the relevant geometric properties of natural rivers (Rinaldo et al., 1992, 2014). Similarly, we adopt simple but hydrologically sound descriptions of habitat size (after Ovaskainen, 2002; Bertassello et al., 2022b) and spatial connectivity (e.g., Bertuzzo et al., 2007; Mari et al., 2014), as well as a prototypical spatially explicit patch occupancy model (Hanski and Ovaskainen, 2000), whose key assumptions have been recently validated (Nicoletti et al., 2023). Our primary objective is to determine the conditions under which a riverine metapopulation bound for asymptotic extinction can temporarily persist through reactive dynamics. Furthermore, we explore whether the temporal scales associated with these transient phenomena may allow for reactive pseudo-persistence (Aiken et al., 2023), wherein recurrent perturbations delay metapopulation extinction indefinitely well below deterministic thresholds.

## Materials and Methods

### Generation of synthetic river networks

We evaluate the asymptotic stability and reactivity of riverine metapopulations in optimal channel networks (OCNs) (Rinaldo et al., 1992, 2014) obtained with the R package OCNet (cran.r-project.org/package=OCNet). The OCN model produces spanning channel networks that are statistically indistinguishable from the real ones extracted from digital terrain models. In particular, OCNs can pass the most stringent similarity tests, such as matching the distribution of total contributing area at a point, which is a more distinctive trait of river networks compared to purely topological metrics (Rinaldo et al., 1998). For this reason, OCNs are widely used to generate stream network analogs for biological and ecological applications (Carraro et al., 2020a; Carraro and Altermatt, 2022). We build the stream network from a square domain made up of 512*×*512 50-m wide pixels, representing a basin with a total surface area of ≈ 655 km^2^. We extract the channel network by imposing that pixels draining areas larger than 8.5 km^2^ are channels. Starting from a pixel description of the channels (Carraro et al., 2020a), we discretize the network into *n* = 51 stream reaches (i.e., the channel segments between two consecutive confluences, or between the head of a first-order stream and the next confluence), for a total length of ≈ 201 km. These streams constitute the fundamental spatial units of the metapopulation model.

### Habitat size

Evaluating the size of a stream in terms of the habitat units it can provide requires making some basic assumptions about the autoecology of the target metapopulation, specifically about how different organisms are influenced by local factors (such as water temperature, nutrient availability, disturbance; Radinger and Wolter, 2015; Boets et al., 2018; Caradima et al., 2021; Negro et al., 2021) and/or behave in the water column (e.g., benthic vs. planktonic or nektonic species; Bertoni, 2011). Environmental factors can be highly site-dependent and, thus, likely characterized by spatial heterogeneities that may not be completely amenable to a general description. Behavioral characteristics, while also highly diversified, can be linked to movement and feeding habits (which also influence the functional role of a species in the context of the lotic community; e.g., Harvey and Altermatt, 2019; Jacquet et al., 2022) that are fundamentally associated with the structure of the river network, which follows basic, almost universal rules (Rodriguez-Iturbe and Rinaldo, 1997).

For example, the hydraulic geometry of stream reaches (Leopold et al., 1964) tells us that, across sections, both the depth and width of the water channel (hence, also their product, that is, the cross-sectional area), for a given discharge with the same frequency of occurrence (say, the mean annual flow in each section), are proportional to such discharge raised to suitable powers (0.4–0.5 for width, 0.3–0.4 for depth; Leopold et al., 1964; Raymond et al., 2012). Furthermore, it can be assumed that the mean annual flow is proportional to the area of the portion of the basin that contributes to the flow section (the landscape-forming discharge is identified by the flood with return period of about 2.2 years, which grants the linear proportionality; Leopold et al., 1964). This total contributing area is thus geometrically linked to both reach depth/width (which may be relevant for organisms living on the riverbed) and cross-sectional area (which may be relevant for floating/swimming organisms). Therefore, we assume that the habitat size *h*_*i*_ of a stream reach (say, *i* ∈ {1, …, *n*}, with *n* being the number of reaches into which the river network has been discretized) is proportional to the product of the total contributing area *A*_*i*_ raised to a suitable power *β* times its channel length *L*_*i*_, i.e., 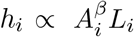 (Ovaskainen, 2002; Muneepeerakul et al., 2008; Bertassello et al., 2022b). For uniform environmental conditions, the values of *β* are expected to range from zero (habitat size independent of the cross-sectional geometry of the river, as in the case of terrestrial organisms living along the river) to one (habitat size dependent on the cross-sectional area; intermediate values ≈ 0.5 would correspond to habitat size dependent on the width/depth of the stream). Values of *β* outside the 0–1 range are also interesting because they could represent the superposition of habitat preferences related, for example, to water temperature (Carraro et al., 2018, 2020b), anthropogenic disturbance (Esselman et al., 2011), or in-stream primary productivity (Segatto et al., 2020), all of which are expected to increase downstream. Therefore, species-specific preferences/tolerances for different environmental conditions could lead to a superlinear increase in habitat size with increasing contributing area (*β >* 1) or even to a decrease (*β <* 0). To allow a fair comparison of metapopulations characterized by different patterns of use of the riverscape as habitat (hence, by different values of *β*), we ultimately define the habitat size of the *i*-th reach as

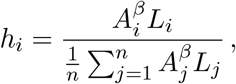

so that any comparison is made using a riverscape with the same (unit) average habitat size. An additional advantage of this “normalization” is that we do not need to assume any arbitrary proportionality constant to link habitat size to geomorphological factors.

### Connectivity

Metapopulation connectivity within the riverscape is described by using a well-established framework (Rinaldo et al., 2020) for the analysis of fluvial networks as ecological corridors (e.g., Bertuzzo et al., 2007; Muneepeerakul et al., 2008; Mari et al., 2011; Carrara et al., 2012; Mari et al., 2012; Ceola et al., 2014; Mari et al., 2014; Bertuzzo et al., 2015; Ciddio et al., 2017; Carraro et al., 2018; Giezendanner et al., 2020; Ma et al., 2020; Bertassello et al., 2022b), in which the dispersal of organisms is represented as a biased random walk over a directed graph where each stream reach is represented by a node. For a given network structure (specifically, for a given connectivity matrix), each inner reach (say, *i*) is characterized by a certain number of closest neighbors downstream (*d*_*i*_; typically, *d*_*i*_ = 1 for all *i*’s except the outlet, for which *d*_*i*_ = 0) and upstream (*u*_*i*_; typically, *u*_*i*_ = 2 for all *i*’s except the headwaters, for which *u*_*i*_ = 0). Let *π* be the probability that the organisms disperse downstream, and let *α* be the probability that the organisms disperse outside the boundaries of the network (assumed to be identical for headwaters and outlets). Other forms of dispersal cost can be accounted for in the description of metapopulation dynamics (Casagrandi and Gatto, 1999). Following Mari et al. (2014), we define the probability *Q*_*ij*_ that an organism disperses from node *i* to *j* as

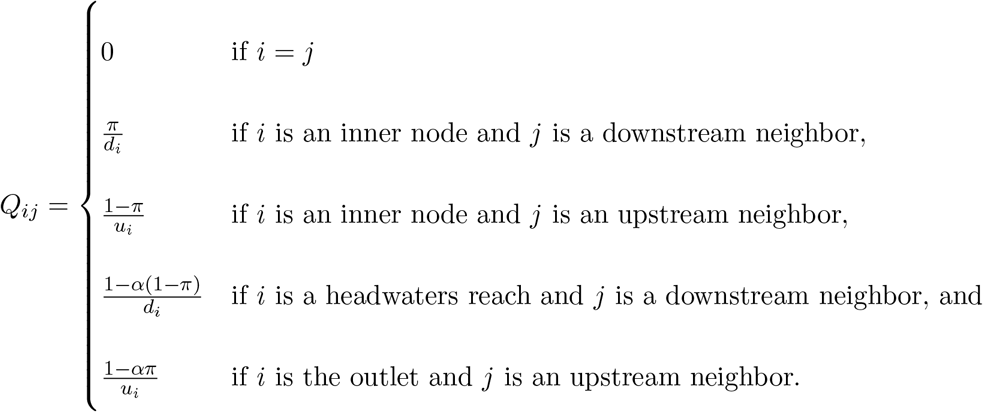

### Patch-occupancy model

Let *p*_*i*_ be the probability that the *i*-th reach (*i* ∈ {1, …, *n*}) of the river network (i.e., the *i*-th patch within the landscape, in the metapopulation jargon) is occupied by a local sub-population. Following Hanski and Ovaskainen (2000), who extended the classic Levins’ (1969) metapopulation model to a spatially explicit framework, the patch-occupancy dynamics can be expressed as the balance between colonization and extinction processes, which can be summarized by the following set of coupled ordinary differential equations (ODEs):

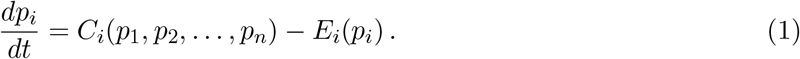

The first term in the right-hand side of the model is the effective colonization rate

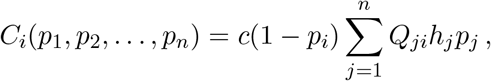

where *c* is the colonization rate (time^−1^) *sensu* Levins (1969), (1 − *p*_*i*_) is the probability that reach *i* is not occupied, and ∑_*j*_ *Q*_*ji*_*h*_*j*_*p*_*j*_ is a measure of the total colonization pressure acting on reach *i* from all other connected patches. The colonization pressure from each patch *j* is evaluated as the product of the probability that an individual disperses from *j* to *i* (*Q*_*ji*_), times the habitat size of reach *j* (*h*_*j*_, which can also be viewed as the simplest proxy of the expected population abundance in *j*), times the probability that reach *j* is indeed occupied (*p*_*j*_). The second term on the right-hand side of model (1), instead, describes local extinction dynamics according to

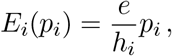

where *e* is the baseline extinction rate (time^−1^), which is modulated by habitat size (following the classical assumption that the smaller the habitat, the larger the extinction probability per unit of time; Hanski, 1999). With proper tuning of the effective colonization (*C*_*i*_) and extinction (*E*_*i*_) rate functions, metapopulation models similar to the one proposed by Hanski and Ovaskainen (2000) have been successfully used to describe patch-occupancy dynamics even in complex, realistic landscapes (see, e.g., Giezendanner et al., 2020; Bertassello et al., 2022a,b; Aiken et al., 2023, for some recent examples).

To reduce the number of parameters, it is possible to measure time in terms of the expected time to extinction in a patch of average habitat size (1*/e*). Introducing the dimensionless quantity *ξ* = *e* · *t*, model (1) thus becomes

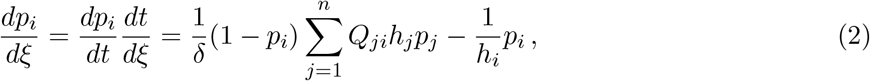

with *δ* = *e/c*. Note that this kind of scaling does not qualitatively alter either the asymptotic stability or the reactivity properties of the model (Lutscher and Wang, 2020).

### Persistence and reactivity

Before proceeding with the analysis of model (2), we briefly recall some basics of asymptotic stability and reactivity analysis that will be used in all the numerical experiments reported in the following. Let

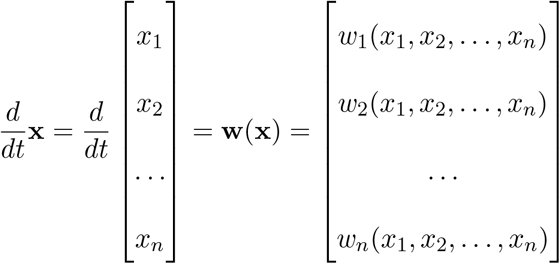

be a system of autonomous ODEs, with **x** and **w**(**x**) being a vector of state variables and the vector field describing their temporal dynamics, respectively.

### Long-term persistence

If an equilibrium 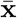 exists, it must satisfy 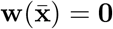, with **0** being the zero column vector of length *n*. The asymptotic stability properties of 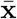 can be (almost always) evaluated by looking at the linearized dynamics of the system, described by the Jacobian matrix 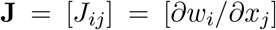 evaluated in a neighborhood of the equilibrium. Specifically, 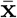 is asymptotically exponentially stable if and only if all the *n* eigenvalues of 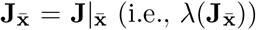 have real parts 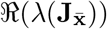 that are all strictly negative, which is equivalent to requiring that

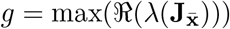

is negative. In this case, all (small) perturbations to 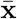 will vanish in the long run, and the system trajectories will eventually converge to the same equilibrium 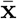.

As an autonomous system (i.e., without external inputs), model (2) always admits an extinction equilibrium 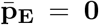 as a trivial solution. Although analytical proof may be non-straightforward, we will show via simulation that an asymptotically stable persistence (i.e., positive) equilibrium appears as soon as the extinction equilibrium loses stability (*g >* 0) because of suitable changes in the underlying model parameterization. We will specifically focus on the asymptotic stability and reactivity properties of 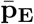, namely we will investigate the ecological conditions conducive to metapopulation reactive/non-reactive extinction or long-term persistence.

### Short-term reactivity

Following Neubert and Caswell (1997), here we define reactivity as the maximum instantaneous growth evaluated with the 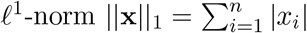 (instead of the *ℓ*^2^-norm 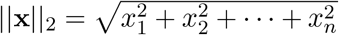 used in the original formulation) achieved by any perturbation departing from an equilibrium point 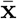, i.e.,

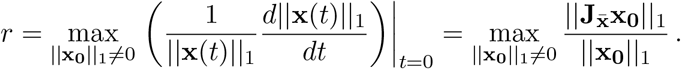

It is straightforward to show (see Harrington et al., 2022, for a lucid demonstration) that, for an equilibrium like 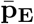 for which only positive perturbations are admissible and for which |*x*_*i*_| = *x*_*i*_, due to the ecological meaning of the variables, reactivity in the *ℓ*^1^-norm corresponds to the maximum element of the column sum (CS) vector of the Jacobian matrix evaluated in a neighborhood of the equilibrium point 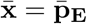, i.e.,

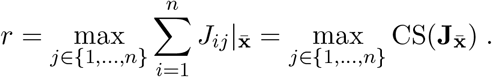

By definition, then, an equilibrium point is termed reactive if *r >* 0. Furthermore, it can be proven (see again Harrington et al., 2022) that the perturbation 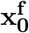 that attains the fastest growth *r* at time *t* = 0 is the one with a single positive component in the *j*^f^-th element of the state vector and otherwise null, with 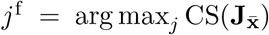. The set *S*^f^ of perturbations (including *j*^f^) for which 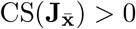 represents the *ℓ*^1^-norm counterpart to the reactivity basin (i.e., the set of initial conditions for which transient growth is possible) that was originally defined using the *ℓ*^2^-norm (Mari et al., 2017).

The growth rate at time *t* = 0 is just one of the features of the so-called amplification envelope (Neubert and Caswell, 1997), which quantifies the maximum growth (again, evaluated here with the *ℓ*^1^-norm) at a generic time *t*^+^ *>* 0 among all possible initial perturbations, that is,

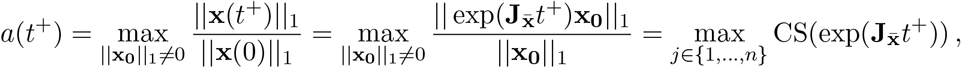

where exp indicates the matrix exponential function. The initial perturbation 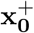 that attains the maximum growth *a*(*t*^+^) at time *t* = *t*^+^ can also be exactly identified (Harrington et al., 2022): specifically, it is the one endowed with a single positive element in the *j*^+^-th element of the state vector and null otherwise, with 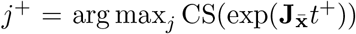. Note that there might exist multiple perturbations for which 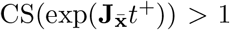, forming a set *S*^+^ (typically different from *S*^f^) of initial conditions characterized by positive growth (*a*(*t*^+^) *>* 1) at a generic time *t*^+^ *>* 0. Among all the 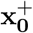 perturbations, of particular interest is 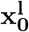, i.e., the one that leads to the maximum growth overall, corresponding to the peak of the amplification envelope occurring at *t*^l^ = arg max_*t*_ *a*(*t*). Note that the computation of the amplification envelope according to the definition given above is based on a linear approximation of the system dynamics, therefore any inference drawn from it will be valid only for small perturbations to the DFE, in non-singular cases where the linearization method holds, and relatively short temporal intervals.

## Results

### Persistence and reactivity conditions using the riverscape matrix

The asymptotic stability properties of the extinction equilibrium 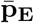 of model (2) are determined by the spectral properties of the Jacobian matrix 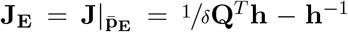, with **Q** = [*Q*_*ij*_] and **h** being a diagonal matrix with positive entries corresponding to habitat size values *h*_*i*_ (*T* indicates matrix transposition). Simple algebraic manipulations show that, in this case, the asymptotic instability condition *g >* 0 can be equivalently written as *ρ*(**M**) *> δ* = *e/c, ρ*(**M**) being the spectral radius of the landscape matrix **M** = **hQ**^*T*^ **h** introduced by Hanski and Ovaskainen (2000), who also referred to *ρ*(**M**) as the metapopulation capacity. Note that *δ* = 1 represents the persistence threshold for Levins’ (1969) metapopulation model (i.e., the spatially implicit version of model (2), for which **M** = 1). Similarly, the reactivity condition *r >* 0 can be rewritten using the landscape matrix as max(CS(**h**) º CS(**h**^−1^**M**)) *> δ*, with º indicating the Hadamard (i.e., element-wise) matrix product.

### Numerical examples of reactive metapopulation dynamics

Clearly, a case of particular interest is that of a metapopulation that is asymptotically headed toward extinction (*g <* 0) but can display reactive behavior in the short term (*r >* 0). For the sake of concreteness, we selected a riparian organism (*β* = 0) characterized by faster extinction than colonization dynamics (*δ* = 2), downstream-biased movement (*π* = 0.8), and costly dispersal (*α* = 0.5), a combination of parameters that leads to asymptotic metapopulation extinction (*g* = −0.033 *<* 0, or *ρ*(**M**) = 1.84 *< δ*, in terms of Hanski and Ovaskainen’s landscape matrix) and transient reactivity (*r* = 1.47 *>* 0 or max(CS(**h**) º CS(**h**^−1^**M**)) = 13.34 *> δ*). Simulation examples of the linearized form of model (2), that is, 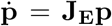, are shown in Figure 1(a, left axis). Individual results for two different initial conditions are reported, corresponding to the perturbations with the fastest initial growth (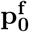, characterized by the maximum growth rate of the state norm at *ξ* = 0; solid blue line) or the largest growth overall (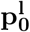, characterized by the maximum growth of the state norm at any time, in this case at *ξ*^l^ = arg max_*ξ*_ *a*(*ξ*) ≈ 5.5 for the linearized form of the model; dashed blue line). Quantitative details aside, these two simulations are characterized by the same qualitative temporal behavior: the state norm (proportional to average patch occupancy) initially grows, decreases, and eventually vanishes. Interestingly, transient dynamics can be quite long-lasting: for instance, in the case of the 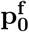 perturbation, the normalized state norm does not go back to the initial size of the perturbation before *ξ* ≈ 26. Ensemble results for all other perturbations leading to a transient amplification of patch occupancy over at least some time scale are also shown (purple shading). For some of these initial conditions, an initial decrease in patch occupancy can indeed be observed, followed by transient increase and asymptotic fade-out. Accounting for density dependence in the simulations of the full model (2) expectedly dampens and shortens the transient growth of the perturbations but does not qualitatively alter the results (Figure S1).

**Figure 1.**
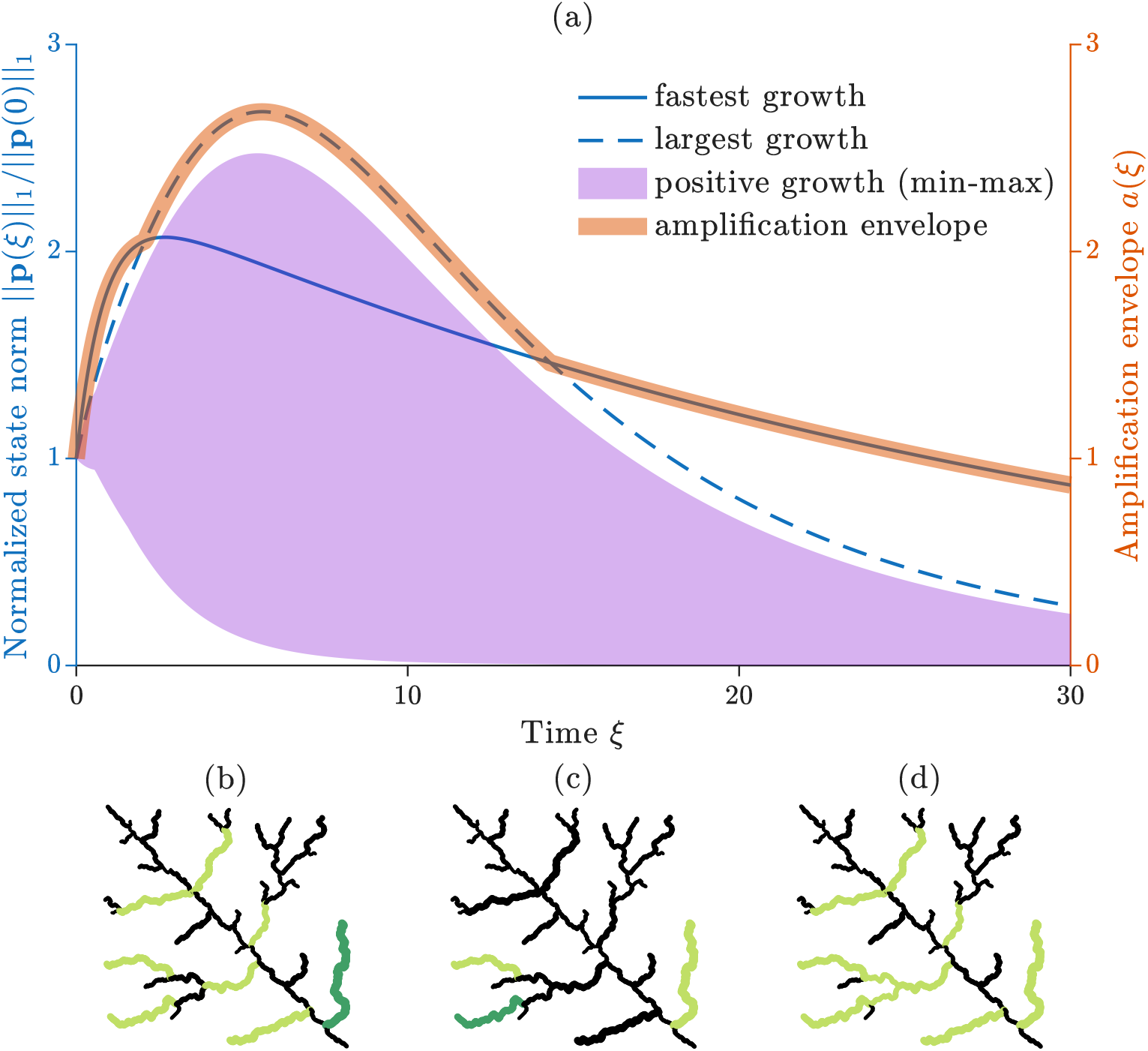
Simulation results for the linearized form of model (2) subject to impulsive perturbations to an asymptotically stable, reactive extinction equilibrium. (a) Temporal dynamics of the normalized *ℓ*^1^-norm of the state vector (proportional to average patch occupancy), as determined by different initial perturbations (blue lines and purple shading, left axis), and amplification envelope (red thick line, right axis). For numerical simulations, initial conditions are set proportionally to the structure of the fastest-growing perturbation (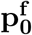, solid blue line), the largest-growing perturbation (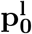, dashed blue lines), or any other perturbation producing a transient growth in patch occupancy over at least some time scale (purple shading). (b) Spatial structure of 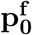 (dark green) and of all single-reach perturbations leading to a positive instantaneous growth rate of patch occupancy at *ξ* = 0 (light green). (c) Spatial structure of 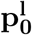 (dark green) and of all single-reach perturbations leading to patch occupancy growth at the peak time *ξ* = *ξ*^l^ of the amplification envelope (light green). (d) Collection of all single-reach perturbations leading to patch occupancy growth over at least some time scales (light green). The analysis is performed on a network with *n* = 51 river reaches, with a single outlet in the bottom-right corner. Parameter values: *δ* = 2, *β* = 0, *π* = 0.8, *α* = 0.5. For this parameter combination, the extinction equilibrium is asymptotically stable (*g* = −0.033) and reactive (*r* = 1.47).

The transient spatiotemporal patterns of patch occupancy generated by the 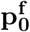 and 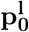 perturbations are illustrated in Figures S2 and S3, respectively. Simulations of model (2) show that, in the case of the fastest-growing perturbation at *ξ* = 0, patch occupancy remains mostly confined to the path that connects the initially colonized stream and the outlet of the network, with the only exception being a spatially limited upstream colonization of the main stem of the river network. In spite of the spatial confinement of patch occupancy, temporal dynamics are quite long-lasting in this case (Figure S2). On the contrary, the perturbation associated with the largest growth overall leads to temporary colonization of a wider intermediate portion of the network, but showing a quicker fading of the probability of patch occupancy (Figure S3).

### Critical perturbations over different time scales

The amplification envelope is also shown in Figure 1(a) (red, right axis). For the selected parameter combination, we find that 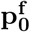 is not only the fastest-growing perturbation at *ξ* = 0 but also the one that grows with the maximum amplitude for *ξ* ⪅ 2 and *ξ* ⪆ 14. Conversely, 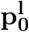 is the transient perturbation that grows the most over the intermediate time interval 2 ⪅ *ξ* ⪅ 14 (which includes *ξ*^l^ ≈ 5.5, the time at which the amplification envelope indeed peaks). The spatial signatures of 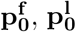, and other perturbations leading to a growth in transient patch-occupancy over some intermediate time scales are shown in panels (b–d) of Figure 1. Specifically, the stream reaches whose initial temporary colonization leads to either the fastest instantaneous growth in patch occupancy (*j*^f^) or the largest growth overall (*j*^l^) are shown in dark green in panels (b) and (c), respectively. In the same panels, the light green reaches are those for which initial colonization leads to an increase in the state norm at *ξ* = 0, albeit at a rate slower than *r* (i.e., those for which 0 *<* CS(**J**_**E**_) *< r*, corresponding to the set 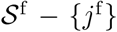; panel b), or at *ξ* = *ξ*^l^, albeit by a factor smaller than *a*(*ξ*^l^) (i.e., those for which 1 *<* CS(exp(**J**_**E**_*ξ*^l^)) *< a*(*ξ*^l^), corresponding to the set *S*^l^ − {*j*^l^}; panel c). All patches whose initial colonization leads to an increase in the state norm over some generic time scale (i.e., those for which CS(exp(**J**_**E**_*ξ*)) *>* 1 for some *ξ*, corresponding to the union of all *S*^*ξ*^) are marked in light green in panel (d). The black reaches in panels (b–d) are instead those for which initial colonization does not lead to a transient growth in patch occupancy over any time scale (i.e., those for which CS(exp(**J**_**E**_*ξ*)) ≤ 1 for all *ξ*’s).

### Reactive pseudo-persistence

The simulation examples reported so far dealt with a metapopulation’s transient response to single, impulsive perturbations. Figure 2 instead shows what may happen when the same asymptotically stable, reactive extinction equilibrium is sporadically but recurrently excited according to different spatial schemes (always assuming single-reach, impulsive perturbations). When perturbations, whose size and timing are shown in panel (a), are randomly localized (‘random seeding’ scenario), only the ones corresponding to the initial colonization of reaches for which CS(exp(**J**_**E**_*ξ*)) *>* 1 for some *ξ* (*S*^*ξ*^, light green reaches in Figure 1(d)) lead to a transient increase in the average probability of patch occupancy within the riverscape over at least some time scale; in all other cases (black reaches in Figure 1(d)), any increase in average occupancy probability is trivially associated with the size of the perturbations. For this reason, when considering an ensemble of multiple realizations of the process, the metapopulation does not display evident reactive behavior (yellow in panel (b); solid line: median, shaded area: min-max range), while still showing a pattern of pseudo-persistence over time, albeit at low average occupancy probabilities. In contrast, if perturbations only insist on reaches belonging to *S*^*ξ*^ (‘positive growth’ scenario), the reactive behavior becomes clearly visible and amplifies the sheer size of the initial perturbations, thus determining marked transient increases in average occupancy probability (purple in panel (b); solid line: median, shaded area: min-max range), which tends to be consistently higher than in the previous case as a result. The same observations apply to cases in which critical perturbations, such as 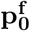 (‘fastest growth’ scenario, solid blue line in panel b) or 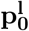 (‘largest growth’ scenario, dashed blue line), repeat over time. However, it is interesting to notice that these perturbations are typically outperformed in terms of average patch occupancy probability by at least some realizations of the ‘positive growth’ scenario, thereby highlighting the contribution of reaches whose colonization leads to a positive yet possibly suboptimal transient growth in average patch occupancy probability in the network. Some examples of patch occupancy patterns generated by model (2) subject to recurrent perturbations are shown in Figures S4–S7.

**Figure 2.**
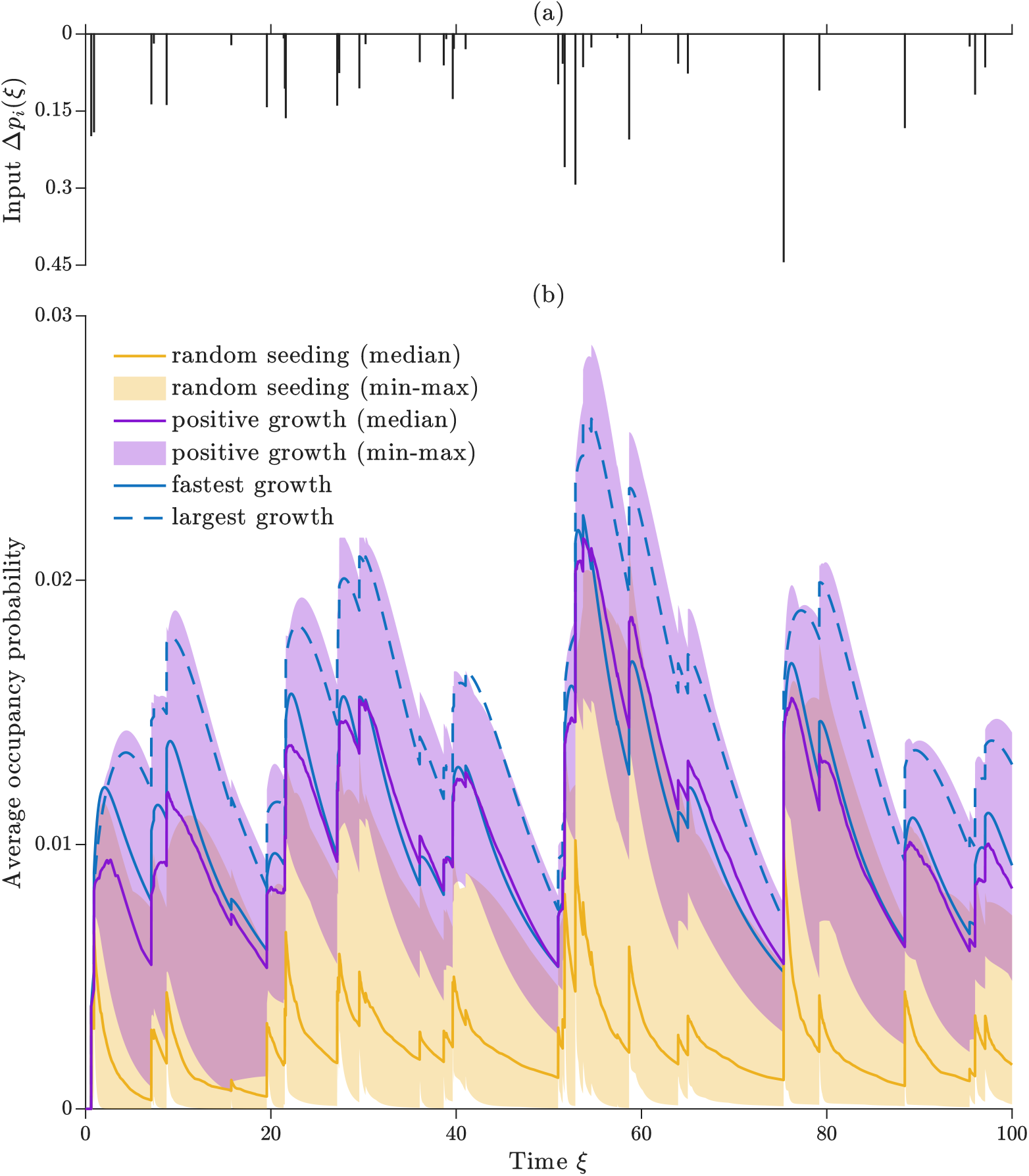
Simulation results for model (2) subject to recurrent impulsive local perturbations to an asymptotically stable, reactive extinction equilibrium. (a) Timing and size of the perturbations. The waiting time between perturbations and the size of each event were randomly generated from exponential distributions with average values of 3 and 0.1, respectively, postulating independence between perturbations (Poisson process). The same sequence of waiting times between events and perturbation sizes was used in each simulation to allow a fair comparison of the various realizations of the process. (b) Trajectories of average patch-occupancy 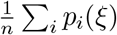 generated by different perturbation scenarios. In the ‘random seeding’ scenario (yellow), the network node *i* to be excited in each perturbation event (*p*_*i*_(*ξ*) = *p*_*i*_(*ξ*) + Δ*p*_*i*_(*ξ*)) was randomly extracted among all network nodes; the curve and the shaded area represent, respectively, the median and the min-max range of average patch occupancy at each time *ξ* obtained considering 100 independent replicas of random seed selection. In the ‘positive growth’ scenario (purple), the network node to be excited in each perturbation was randomly selected among the network nodes with CS(exp(**J**_**E**_*ξ*)) *>* 1 for some *ξ* after perturbation (green reaches in Figure 1(d)); other details as in the previous scenario. In the ‘fastest growth’ scenario (solid blue line), the perturbation with the fastest instantaneous amplification 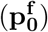 was always applied. In the ‘largest growth’ scenario (dashed blue line), the perturbation with the largest amplification overall 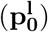 was always applied. Parameters and other details as in Figure 1.

### Parametric conditions for persistence and reactivity

In line with the Levins-like origins of model (2), a key parameter determining the asymptotic stability and reactivity properties of the extinction equilibrium of model (2) is *δ* = *e/c*, that is, the extinction-to-colonizat rate ratio of the original model. As shown in Figure 3, starting from the stable and reactive steady state shown in Figure 1, increasing values of *δ* lead to decreasing (but still positive) values of *r* and, eventually, to non-reactive extinction equilibria (*r <* 0). In contrast, decreasing values of *δ* lead to increasing values of *g*. For values of *δ* ⪅ 2, the extinction equilibrium loses stability (*g >* 0) and an asymptotically stable persistence equilibrium (i.e., endowed with at least one strictly positive component) appears. Long-term patch occupancy is obviously null with *g <* 0 (independently of the sign of *r*), while it increases quickly for decreasing values of *δ* below the asymptotic stability threshold.

**Figure 3.**
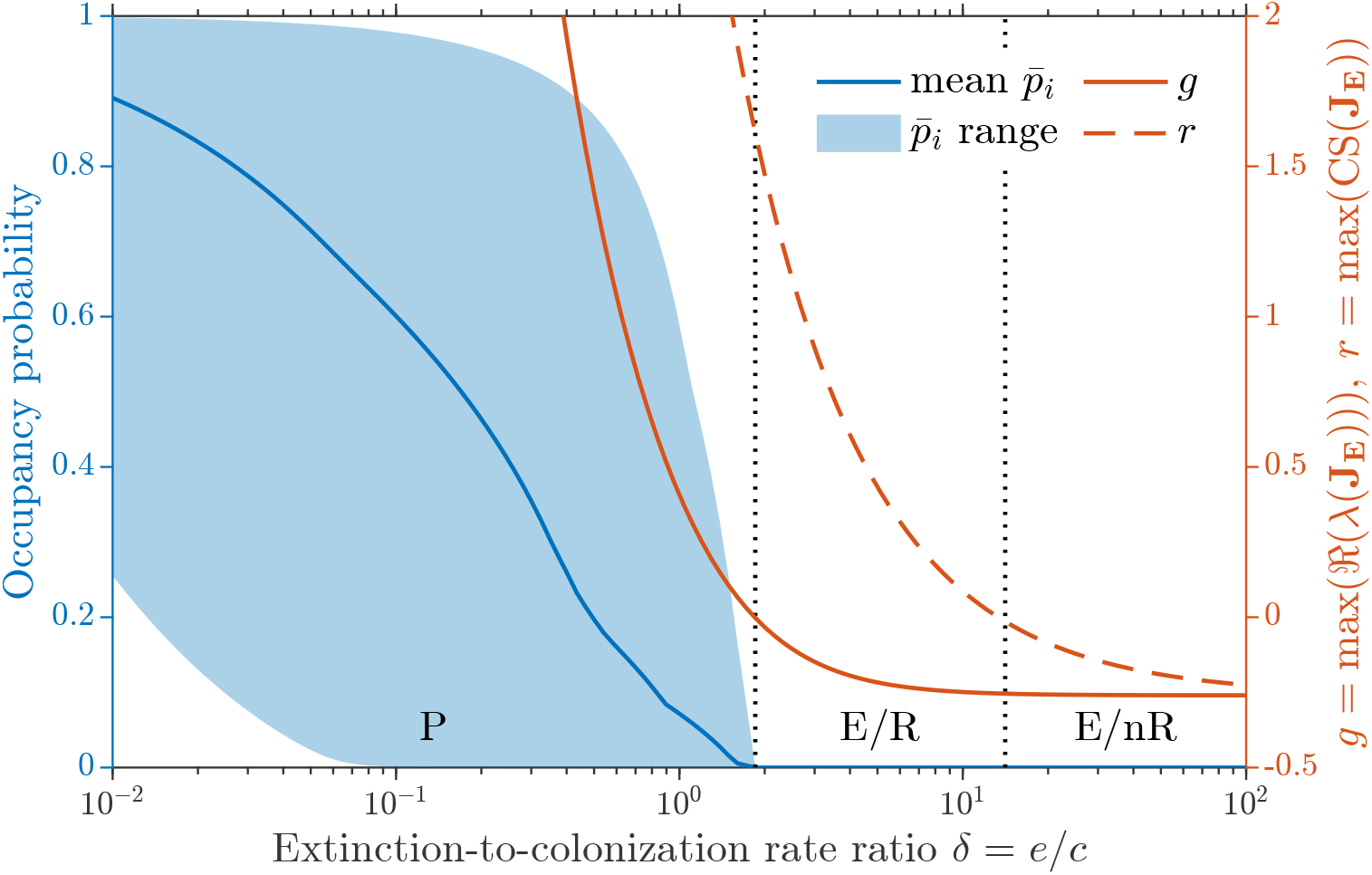
Asymptotic stability and reactivity properties of the equilibria of model (2) as a function of the extinction-to-colonization ratio *δ*. The long-term average patch occupancy 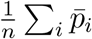 is shown as a blue line, while the min-max range of the steady-state reach occupancy probabilities is shown as a blue shading (left axis). The quantity *g* = max(ℜ(*λ*(**J**_**E**_))) is shown as a red solid line, while *r* = max CS(**J**_**E**_) is shown as a red dashed line (right axis). The vertical dotted lines separate parametric regions of species persistence (abbreviated as P) from parametric regions in which the extinction equilibrium is asymptotically stable and reactive (E/R) or asymptotically stable and non-reactive (E/nR). Parameters (except *δ*) and other details as in Figure 1.

A more comprehensive catalog of the asymptotic stability and reactivity properties of the extinction equilibrium of system (2) as a function of the model parameters is shown in Figure 4. Some interesting results emerge with respect to the extinction-to-colonization rate ratio and the area-suitability exponent (panel (a)). Reactive behavior (and possibly reactive persistence) is only found for intermediate values of *δ*, as already observed in Figure 3. Interestingly, though, the range of *δ* values (between the solid and dashed lines, representing the conditions *g* = 0 and *r* = 0, respectively) conducive to an asymptotically stable, reactive metapopulation extinction equilibrium expands sensibly for *β* ⪅ 0.5. This result is structurally robust to changes in the parameterization of dispersal and is dramatically amplified by an increase in the bias of downstream movement (large *π*; Figure S8).

**Figure 4.**
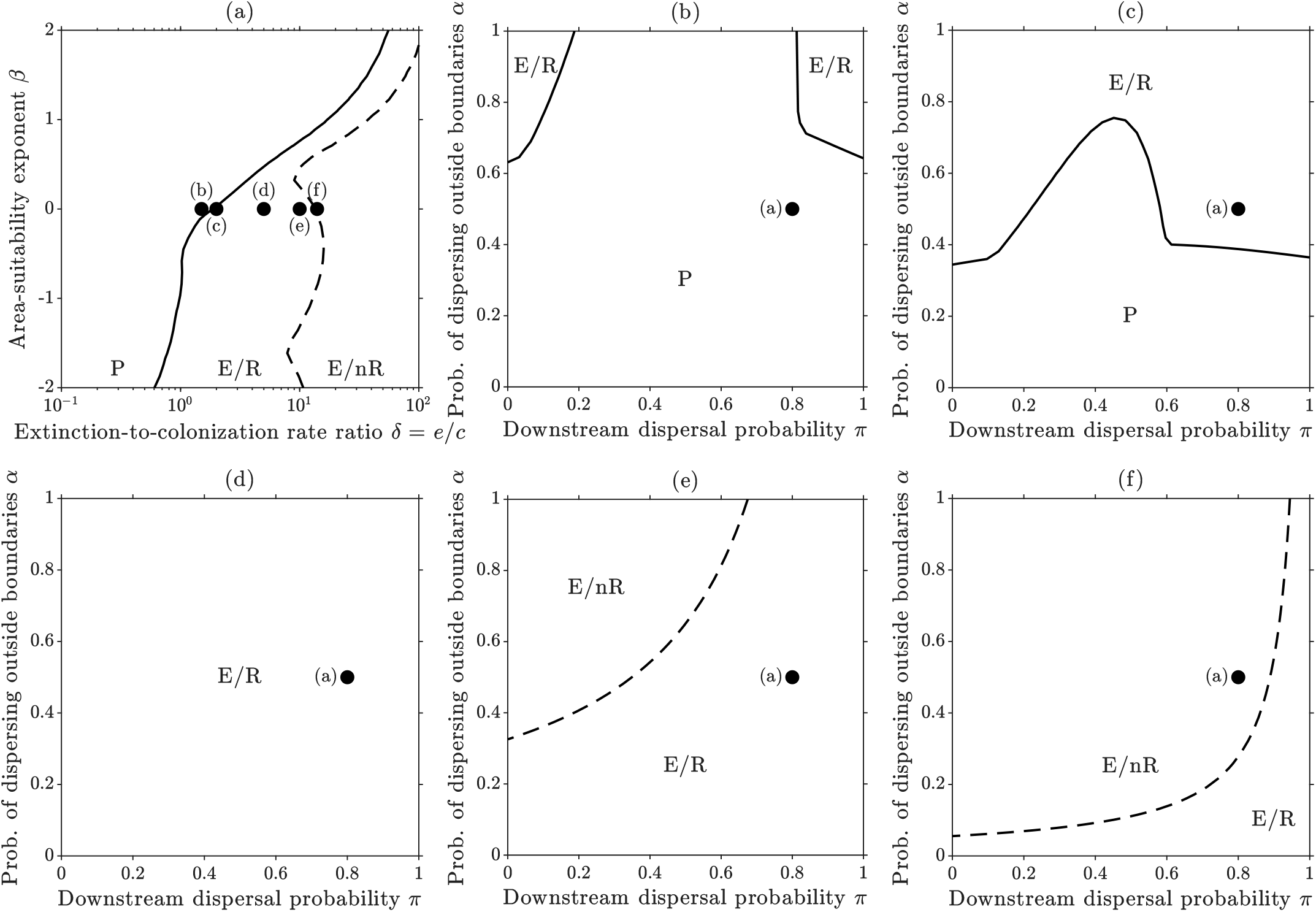
A catalog of the asymptotic stability and reactivity properties of the extinction equilibrium of model (2). (a) Stability and reactivity regions in the parameter space (*δ, β*), spanning the extinction-to-colonization rate ratio and the area-suitability exponent. (b–f) Stability and reactivity regions in the parameter space (*π, α*), spanning the downstream dispersal probability and the probability of dispersing outside the habitat boundaries. In each panel, the solid curve identifies the parameter combinations for which *g* = 0 (or *ρ*(**M** = *δ*)), which separates the regions (marked as E) in which the extinction equilibrium is asymptotically stable (and the metapopulation cannot persist) from those (marked as P) in which the equilibrium is asymptotically unstable (and the metapopulation can persist). The dashed curve, instead, identifies the condition *r* = 0 (or max(CS(**h**) º CS(**h**^−1^**M**)) = *δ*), which separates the regions (marked as R) where the asymptotically stable extinction equilibrium is reactive (and transient increases in patch occupancy can occur) from those (marked with nR) in which the equilibrium is non-reactive (and transient increases in patch occupancy cannot occur). The dots in panel (a) show the parameter combinations explored in panels (b–f), while the dots in panels (e–f) refer to the parameterization of panel (a). Note that no curves are shown in panel (d) because for that combination of *δ* and *β* it is not possible to alter the stability and reactivity properties of the extinction equilibrium by changing only the dispersal parameters *π* and *α*. Other details as in Figure 1.

Regarding dispersal parameters (Figure 4, panels (b–f)), their impact on the asymptotic stability and reactivity properties of the extinction equilibrium can be better observed close to the stability/reactivity boundaries, as changes in the parameterization of dispersal alone may not be sufficient to alter the stability/reactivity properties of the extinction equilibrium (panel (d)). The asymptotic stability of the extinction equilibrium (above the solid lines, which are only visible for the values of *δ* considered in panels (b) and (c)) is favored by high values of the probability of dispersal outside the limits of the river network (large *α*), especially for strongly biased dispersal (either small or large *π*). Reactive behavior (below the dashed lines, which are only visible for the values of *δ* considered in panels (e) and (f)) is favored instead by a low probability of dispersal outside the boundaries of the river network (small *α*) and downstream-biased dispersal (large *π*). Changes in the area-suitability exponent do not qualitatively alter this picture (Figures S9).

## Discussion

In this study, we have proposed a modeling framework to investigate transient pseudo-persistence in riverine metapopulation dynamics, i.e., the temporary occupation of riverscape patches by a metapopulation that is bound to go extinct in the long run (Aiken et al., 2023). The theoretical foundation of our approach is rooted in the concept of ecological reactivity (Neubert and Caswell, 1997), which provides an effective complement to asymptotic stability analysis for the study of metapopulation persistence on short to long time scales (Mari et al., 2017). Here, we have elaborated on it specifically focusing on fluvial landscapes, which often support high levels of biodiversity (Dudgeon et al., 2006), but may be particularly vulnerable to disturbance and environmental change (Reid et al., 2019). The use of synthetic, yet realistic, riverscapes (Rinaldo et al., 2014; Carraro et al., 2020a), coupled with comprehensive descriptions of local habitat suitability (Ovaskainen, 2002; Bertassello et al., 2022b) and dispersal along the stream (Bertuzzo et al., 2007; Mari et al., 2014), allowed us to abstract our results from a specific case study and search for general ecological conditions conducive to transient patch occupancy.

Our results indicate that, under suitable ecohydrological conditions, even metapopulations that are bound to extinction in the long run can colonize parts of a riverscape for non-negligible periods of time. For example, we found that a single perturbation (such as the import of a few colonizing individuals from an external reservoir) to an asymptotically stable extinction equilibrium can lead to transient dynamics with a relaxation period more than 20 times longer than the expected local persistence time (Figure 1). Under these conditions, repeated perturbations can delay the eventual deterministic extinction of the metapopulation almost indefinitely (Figure 2), thus creating a transient pattern of reactive pseudo-persistence (Aiken et al., 2023). Although this extinction debt may provide a window of opportunity for conservation, delayed extinction also poses a significant challenge, especially in terms of the prolonged monitoring effort that might be required to distinguish between short- and long-term metapopulation recovery (Kuussaari et al., 2009). Our findings also lend support to the practice of periodic reintroductions in ecological restoration efforts, not only to overcome biotic homogenization (Holl et al., 2022) but also to “buy time” for struggling metapopulations and favor their (transient) persistence. In this sense, reactivity analysis can help identify habitat patches that are expected to lead to the largest transient amplifications on different temporal scales.

A particularly instructive empirical illustration of the dynamics described by our model is provided by the long-term study of marble trout (*Salmo marmoratus*) populations in the highly fragmented river networks of western Slovenia. These isolated populations, confined to small headwater reaches separated by natural and anthropogenic barriers, are subject to recurrent flash floods and debris flows that episodically reduce local abundances to near-zero. Demographic monitoring and genetic analyses have documented severe bottlenecks followed by partial recovery (Vincenzi et al., 2016, 2017), a pattern consistent with transient reactive dynamics in a metapopulation operating close to or below its asymptotic extinction threshold. The conservation program for this species, which has relied on periodic translocation of individuals into previously unoccupied or depopulated headwater reaches since 1993 (Vincenzi et al., 2012), further mirrors the recurrent reintroduction scenarios explored in our study: even when individual introduction events are small, their repeated application to reaches with favorable reactive properties may substantially delay the decline of the metapopulation. Our framework thus suggests that spatial targeting of such translocation efforts, prioritizing reaches whose colonization is expected to produce the largest transient amplification of patch occupancy, could improve the effectiveness of restoration programs for riverine salmonids and other fragmented freshwater species facing comparable threats.

Our analysis also showed that reactive behavior can only be found for intermediate values of the extinction-to-colonization rate ratio *δ* (Figure 3), with the lower bound given by the asymptotic stability threshold. Interestingly, this threshold does not necessarily correspond to *δ* = 1 as in the case of spatially implicit metapopulation models (Levins, 1969), as already shown by previous research on spatially explicit metapopulations (Hanski and Ovaskainen, 2000) also in the context of fluvial landscapes (Mari et al., 2014). The upper limit of reactive behavior can correspond to values of *δ* that are much higher than the extinction threshold (in fact, close to or even greater than 10 for most of the parameter combinations tested; Figure 4a). This finding suggests that reactive persistence may be quite common for many species close to their extinction threshold. The area-suitability exponent *β* also proved to be an important factor, with reactive behavior more common (or at least occurring for wider ranges of the extinction-to-colonization rate ratio) for values of *β* below approximately 0.5 (Figure 4a) in the presence of downstream-biased dispersal. From an ecohydrological perspective, this may suggest that benthic species (whose habitat size, for a given river section, depends on the width of the stream) are more likely to exhibit reactive behavior than planktonic or nektonic species (whose habitat size depends on the cross-sectional area). Species that are particularly suited to live in the headwaters (hence *β <* 0), such as shredders feeding on coarse particulate organic material that is more abundant in the upper reaches of the river network (Vannote et al., 1980), would be expected to show the strongest reactive behavior.

We found that the dispersal parameters also play a role in determining the possible appearance of reactive behavior, although only relatively close to the stability/reactivity boundaries (Figure 4b). In particular, near the asymptotic extinction boundary, reactive behavior is only possible if dispersal is sufficiently costly (large values of *α*) and biased (small or large *π*). This result echoes previous research linking reactivity in metapopulation dynamics to dispersal asymmetries (e.g., as determined by ocean currents, as in Aiken and Navarrete, 2011). The ecological explanation for this phenomenon is that, in the presence of asymmetric dispersal, small positive perturbations to a patch that is a transient ‘source’ of dispersing propagules within the metapopulation (Harrington et al., 2022) can cause a temporary increase in patch occupancy at the landscape scale (Aiken et al., 2023). Mathematically, asymmetry makes the eigenvectors of the Jacobian matrix of the system evaluated in a neighborhood of the extinction equilibrium non-orthogonal, which is a necessary (but not sufficient) condition for reactivity (Farrell and Ioannou, 1996). Further away from the asymptotic extinction threshold, reactive behavior instead requires a sufficiently low cost of dispersal and/or a sufficiently strong downstream bias in dispersal. This observation suggests that the role of dispersal in the occurrence of reactive behavior should always be assessed in conjunction with that of the other processes that govern the dynamics of the metapopulation. We have previously shown that the superposition of multiple dispersal routes (such as in the case of amphibian species that can disperse both along the river and overland) can promote the asymptotic persistence of riverine metapopulations (Mari et al., 2014). Assessing whether and how multi-modal dispersal can influence reactive behavior and, possibly, pseudo-persistence may prove a worthwhile direction for future research.

Here, we have focused on the asymptotic stability and reactivity properties of the metapopulation extinction equilibrium. The motivation for our choice is twofold. First, from an ecological perspective, understanding the conditions for the short- and long-term persistence of a riverine (meta)population is a crucial challenge for conservation. Second, from a technical point of view, the study of equilibria endowed with strictly positive components (as expected in the case of a non-trivial equilibrium solution of model (2)) may require an extension of the operational definition of reactivity because, in that case, a reactive deviation from the equilibrium point would not necessarily be characterized by positive components (Harrington et al., 2022). Technical challenges notwithstanding, we believe that studying the reactivity properties of asymptotically persistent riverine metapopulations may be another compelling avenue for future research. Another interesting direction for future research may be represented by the analysis of the reactive behavior of metapopulations living in dynamic (and possibly ephemeral) fluvial environments, where the determinants of the landscape matrix (local habitat suitability and spatial connectivity) or even the topological configuration of the network can change over time (Giezendanner et al., 2021; Bertassello et al., 2022a,b; Durighetto et al., 2023), which will require an extension of the definition of ecological reactivity to time-varying systems (see Vesipa and Ridolfi, 2017; Lutscher and Wang, 2020, for some notable precedents).

In conclusion, our work has shown that, under suitable ecohydrological conditions, reactivity can favor the transient persistence of riverine metapopulations. Such pseudo-persistence can occur well below the extinction threshold and, in the case of recurrent colonization events from an external source, even for arbitrarily long periods of time. Our approach thus offers a broader perspective for the study of metapopulation dynamics, one that complements classical stability analysis and relaxes the stricter requirements of asymptotic persistence. Although our study is mainly theoretical, it is not devoid of practical implications. Indeed, our results can help in the evaluation of different conservation strategies aimed at improving the likelihood of metapopulation persistence over different time scales and guide the selection of the most promising sites for restoration efforts.

## Acknowledgments

L.M. and R.C. acknowledge support from the National Recovery and Resilience Plan (NRRP), Mission 4 Component 2 Investment 1.4 - Call for tender No. 3138 of 16 December 2021, rectified by Decree n.3175 of 18 December 2021 of Italian Ministry of University and Research funded by the European Union – NextGenerationEU; Award Number: Project code CN 00000033, Concession Decree No. 1034 of 17 June 2022 adopted by the Italian Ministry of University and Research, CUP H43C22000530, Project title “National Biodiversity Future Center - NBFC”. A.R. acknowledges funding from the Swiss NSF via the project “Optimal control of intervention strategies for waterborne disease epidemics” (200021-172578).

## Data and code availability

The data and the main code needed to perform the analyses described in this work will be available at https://github.com/lorenzo-mari/reactivity-river-networks.

## References

Aiken, C. M. and Navarrete, S. A. (2011). Environmental fluctuations and asymmetrical dispersal: generalized stability theory for studying metapopulation persistence and marine protected areas. Marine Ecology Progress Series, 428:77–88.

Aiken, C. M., Navarrete, S. A., and Jackson, E. L. (2023). Reactive persistence, spatial management, and conservation of metapopulations: An application to seagrass restoration. Ecological Applications, 33(2):e2774.

Altermatt, F. (2013). Diversity in riverine metacommunities: a network perspective. Aquatic Ecology, 47:365–377.

Belletti, B., Garcia de Leaniz, C., Jones, J., Bizzi, S., Börger, L., Segura, G., Castelletti, A., Van de Bund, W., Aarestrup, K., Barry, J., et al. (2020). More than one million barriers fragment europe’s rivers. Nature, 588(7838):436–441.

Bertassello, L., Bertuzzo, E., Botter, G., Jawitz, J., Aubeneau, A., Hoverman, J., Rinaldo, A., and Rao, P. (2021). Dynamic spatio-temporal patterns of metapopulation occupancy in patchy habitats. Royal Society Open Science, 8(1):201309.

Bertassello, L., Jawitz, J., Bertuzzo, E., Botter, G., Rinaldo, A., Aubeneau, A., Hoverman, J., and Rao, P. (2022a). Persistence of amphibian metapopulation occupancy in dynamic wetlandscapes. Landscape Ecology, 37(3):695–711.

Bertassello, L. E., Durighetto, N., and Botter, G. (2022b). Eco-hydrological modelling of channel network dynamics—Part 2: Application to metapopulation dynamics. Royal Society Open Science, 9(11):220945.

Bertoni, R. (2011). Limnology of rivers and lakes. In Gobal, B., editor, Encyclopedia of Life Support Systems (EOLSS), chapter Limnology, pages 1–68. Developed under the Auspices of the UNESCO, EOLSS Publishers, Oxford, UK. http://www.eolss.net.

Bertuzzo, E., Maritan, A., Gatto, M., Rodriguez-Iturbe, I., and Rinaldo, A. (2007). River networks and ecological corridors: Reactive transport on fractals, migration fronts, hydrochory. Water Resources Research, 43(4).

Bertuzzo, E., Rodriguez-Iturbe, I., and Rinaldo, A. (2015). Metapopulation capacity of evolving fluvial landscapes. Water Resources Research, 51(4):2696–2706.

Boets, P., Gobeyn, S., Dillen, A., Poelman, E., and Goethals, P. L. (2018). Assessing the suitable habitat for reintroduction of brown trout (Salmo trutta forma fario) in a lowland river: A modeling approach. Ecology and Evolution, 8(10):5191–5205.

Caradima, B., Scheidegger, A., Brodersen, J., and Schuwirth, N. (2021). Bridging mechanistic conceptual models and statistical species distribution models of riverine fish. Ecological Modelling, 457:109680.

Carrara, F., Altermatt, F., Rodriguez-Iturbe, I., and Rinaldo, A. (2012). Dendritic connectivity controls biodiversity patterns in experimental metacommunities. Proceedings of the National Academy of Sciences, 109(15):5761–5766.

Carraro, L. and Altermatt, F. (2022). Optimal channel networks accurately model ecologically-relevant geomorphological features of branching river networks. Communications Earth & Environment, 3(1):125.

Carraro, L., Bertuzzo, E., Fronhofer, E. A., Furrer, R., Gounand, I., Rinaldo, A., and Altermatt, F. (2020a). Generation and application of river network analogues for use in ecology and evolution. Ecology and Evolution, 10(14):7537–7550.

Carraro, L., Mari, L., Gatto, M., Rinaldo, A., and Bertuzzo, E. (2018). Spread of proliferative kidney disease in fish along stream networks: A spatial metacommunity framework. Freshwater Biology, 63(1):114–127.

Carraro, L., Toffolon, M., Rinaldo, A., and Bertuzzo, E. (2020b). SESTET: A spatially explicit stream temperature model based on equilibrium temperature. Hydrological Processes, 34(2):355–369.

Casagrandi, R. and Gatto, M. (1999). A mesoscale approach to extinction risk in fragmented habitats. Nature, 400(6744):560–562.

Ceola, S., Bertuzzo, E., Singer, G., Battin, T. J., Montanari, A., and Rinaldo, A. (2014). Hydrologic controls on basin-scale distribution of benthic invertebrates. Water Resources Research, 50(4):2903–2920.

Ciddio, M., Mari, L., Sokolow, S. H., De Leo, G. A., Casagrandi, R., and Gatto, M. (2017). The spatial spread of schistosomiasis: A multidimensional network model applied to saint-louis region, senegal. Advances in water resources, 108:406–415.

Dudgeon, D., Arthington, A. H., Gessner, M. O., Kawabata, Z.-I., Knowler, D. J., Lévêque, C., Naiman, R. J., Prieur-Richard, A.-H., Soto, D., Stiassny, M. L., et al. (2006). Freshwater biodiversity: Importance, threats, status and conservation challenges. Biological Reviews, 81(2):163–182.

Durighetto, N., Noto, S., Tauro, F., Grimaldi, S., and Botter, G. (2023). Integrating spatially-and temporally-heterogeneous data on river network dynamics using graph theory. Iscience, 26(8).

Esselman, P. C., Infante, D. M., Wang, L., Wu, D., Cooper, A. R., and Taylor, W. W. (2011). An index of cumulative disturbance to river fish habitats of the conterminous United States from landscape anthropogenic activities. Ecological Restoration, pages 133–151.

Fagan, W. F. (2002). Connectivity, fragmentation, and extinction risk in dendritic metapopulations. Ecology, 83(12):3243–3249.

Farrell, B. F. and Ioannou, P. J. (1996). Generalized stability theory. Part I: Autonomous operators. Journal of Atmospheric Sciences, 53(14):2025–2040.

Fuller, M. R., Doyle, M. W., and Strayer, D. L. (2015). Causes and consequences of habitat fragmentation in river networks. Annals of the New York Academy of Sciences, 1355(1):31–51.

Giezendanner, J., Benettin, P., Durighetto, N., Botter, G., and Rinaldo, A. (2021). A note on the role of seasonal expansions and contractions of the flowing fluvial network on metapopulation persistence. Water Resources Research, 57(11):e2021WR029813.

Giezendanner, J., Pasetto, D., Perez-Saez, J., Cerrato, C., Viterbi, R., Terzago, S., Palazzi, E., and Rinaldo, A. (2020). Earth and field observations underpin metapopulation dynamics in complex landscapes: Near-term study on carabids. Proceedings of the National Academy of Sciences, 117(23):12877–12884.

Haase, P., Bowler, D. E., Baker, N. J., Bonada, N., Domisch, S., Garcia Marquez, J. R., Heino, J., Hering, D., Jähnig, S. C., Schmidt-Kloiber, A., et al. (2023). The recovery of european freshwater biodiversity has come to a halt. Nature, pages 1–7.

Hanski, I. (1999). Metapopulation Ecology. Oxford University Press.

Hanski, I. and Ovaskainen, O. (2000). The metapopulation capacity of a fragmented landscape. Nature, 404(6779):755–758.

Hanski, I. and Ovaskainen, O. (2002). Extinction debt at extinction threshold. Conservation Biology, 16(3):666–673.

Harrington, P. D., Lewis, M. A., and van den Driessche, P. (2022). Reactivity, attenuation, and transients in metapopulations. SIAM Journal on Applied Dynamical Systems, 21(2):1287–1321.

Harvey, E. and Altermatt, F. (2019). Regulation of the functional structure of aquatic communities across spatial scales in a major river network. Ecology, 100(4):e02633.

Hastings, A. (2004). Transients: the key to long-term ecological understanding? Trends in Ecology & Evolution, 19(1):39–45.

Holl, K. D., Luong, J. C., and Brancalion, P. H. (2022). Overcoming biotic homogenization in ecological restoration. Trends in Ecology & Evolution.

Huang, Q. and Lewis, M. A. (2015). Homing fidelity and reproductive rate for migratory populations. Theoretical Ecology, 8:187–205.

Jacquet, C., Carraro, L., and Altermatt, F. (2022). Meta-ecosystem dynamics drive the spatial distribution of functional groups in river networks. Oikos, 2022(11):e09372.

Kuussaari, M., Bommarco, R., Heikkinen, R. K., Helm, A., Krauss, J., Lindborg, R., Öckinger, E., Pärtel, M., Pino, J., Rodà, F., et al. (2009). Extinction debt: A challenge for biodiversity conservation. Trends in Ecology & Evolution, 24(10):564–571.

Leopold, L. B., Wolman, M. G., and Miller, J. P. (1964). Fluvial processes in geomorphology. Freeman.

Levins, R. (1969). Some demographic and genetic consequences of environmental heterogeneity for biological control. Bulletin of the ESA, 15(3):237–240.

Lutscher, F., Nisbet, R. M., and Pachepsky, E. (2010). Population persistence in the face of advection. Theoretical Ecology, 3:271–284.

Lutscher, F. and Wang, X. (2020). Reactivity of communities at equilibrium and periodic orbits. Journal of Theoretical Biology, 493:110240.

Ma, C., Shen, Y., Bearup, D., Fagan, W. F., and Liao, J. (2020). Spatial variation in branch size promotes metapopulation persistence in dendritic river networks. Freshwater Biology, 65(3):426–434.

MacArthur, R. H. and Wilson, E. O. (1967). The theory of island biogeography. Princeton University Press.

Mari, L., Bertuzzo, E., Casagrandi, R., Gatto, M., Levin, S. A., Rodriguez-Iturbe, I., and Rinaldo, A. (2011). Hydrologic controls and anthropogenic drivers of the zebra mussel invasion of the Mississippi-Missouri river system. Water Resources Research, 47(3).

Mari, L., Bertuzzo, E., Righetto, L., Casagrandi, R., Gatto, M., Rodriguez-Iturbe, I., and Rinaldo, A. (2012). Modelling cholera epidemics: The role of waterways, human mobility and sanitation. Journal of the Royal Society interface, 9(67):376–388.

Mari, L., Casagrandi, R., Bertuzzo, E., Rinaldo, A., and Gatto, M. (2014). Metapopulation persistence and species spread in river networks. Ecology Letters, 17(4):426–434.

Mari, L., Casagrandi, R., Rinaldo, A., and Gatto, M. (2017). A generalized definition of reactivity for ecological systems and the problem of transient species dynamics. Methods in Ecology and Evolution, 8(11):1574–1584.

Mari, L., Trevisin, C., Rinaldo, A., and Gatto, M. (2025). Sufficient reproduction numbers to prevent recurrent epidemics. Methods in Ecology and Evolution. In press.

Matessi, C. and Gatto, M. (1984). Does K-selection imply prudent predation? Theoretical Population Biology, 25(3):347–363.

Muneepeerakul, R., Bertuzzo, E., Lynch, H. J., Fagan, W. F., Rinaldo, A., and Rodriguez-Iturbe, I. (2008). Neutral metacommunity models predict fish diversity patterns in Mississippi–Missouri basin. Nature, 453(7192):220–222.

Negro, G., Fenoglio, S., Quaranta, E., Comoglio, C., Garzia, I., and Vezza, P. (2021). Habitat preferences of italian freshwater fish: A systematic review of data availability for applications of the MesoHABSIM model. Frontiers in Environmental Science, 9:634737.

Neubert, M. G. and Caswell, H. (1997). Alternatives to resilience for measuring the responses of ecological systems to perturbations. Ecology, 78(3):653–665.

Nguyen, T. D., Tang, T., Veprauskas, A., Wu, Y., and Zhou, Y. (2026). Impact of network connectivity on the dynamics of populations in stream environments. Journal of Mathematical Biology, 92(4):47.

Nicoletti, G., Padmanabha, P., Azaele, S., Suweis, S., Rinaldo, A., and Maritan, A. (2023). Emergent encoding of dispersal network topologies in spatial metapopulation models. Proceedings of the National Academy of Sciences, 120(46):e2311548120.

Ovaskainen, O. (2002). Long-term persistence of species and the SLOSS problem. Journal of Theoretical Biology, 218(4):419–433.

Pachepsky, E., Lutscher, F., Nisbet, R., and Lewis, M. A. (2005). Persistence, spread and the drift paradox. Theoretical Population Biology, 67(1):61–73.

Petsch, D. K., Cionek, V. d. M., Thomaz, S. M., and dos Santos, N. C. L. (2023). Ecosystem services provided by river-floodplain ecosystems. Hydrobiologia, 850(12–13):2563–2584.

Radinger, J. and Wolter, C. (2015). Disentangling the effects of habitat suitability, dispersal, and fragmentation on the distribution of river fishes. Ecological Applications, 25(4):914–927.

Raymond, P. A., Zappa, C. J., Butman, D., Bott, T. L., Potter, J., Mulholland, P., Laursen, A. E., McDowell, W. H., and Newbold, D. (2012). Scaling the gas transfer velocity and hydraulic geometry in streams and small rivers. Limnology and Oceanography: Fluids and Environments, 2(1):41–53.

Reid, A. J., Carlson, A. K., Creed, I. F., Eliason, E. J., Gell, P. A., Johnson, P. T., Kidd, K. A., MacCormack, T. J., Olden, J. D., Ormerod, S. J., et al. (2019). Emerging threats and persistent conservation challenges for freshwater biodiversity. Biological Reviews, 94(3):849–873.

Riis, T., Kelly-Quinn, M., Aguiar, F. C., Manolaki, P., Bruno, D., Bejarano, M. D., Clerici, N., Fernandes, M. R., Franco, J. C., Pettit, N., et al. (2020). Global overview of ecosystem services provided by riparian vegetation. BioScience, 70(6):501–514.

Rinaldo, A., Gatto, M., and Rodriguez-Iturbe, I. (2020). River networks as ecological corridors: Species, populations, pathogens. Cambridge University Press.

Rinaldo, A., Rigon, R., Banavar, J. R., Maritan, A., and Rodriguez-Iturbe, I. (2014). Evolution and selection of river networks: Statics, dynamics, and complexity. Proceedings of the National Academy of Sciences, 111(7):2417–2424.

Rinaldo, A., Rodriguez-Iturbe, I., and Rigon, R. (1998). Channel networks. Annual Review of Earth and Planetary Sciences, 26(1):289–327.

Rinaldo, A., Rodriguez-Iturbe, I., Rigon, R., Bras, R. L., Ijjasz-Vasquez, E., and Marani, A. (1992). Minimum energy and fractal structures of drainage networks. Water Resources Research, 28(9):2183–2195.

Rodriguez-Iturbe, I. and Rinaldo, A. (1997). Fractal river basins: Chance and self-organization. Cambridge University Press.

Roughgarden, J. (1979). Theory of population genetics and evolutionary ecology: An introduction. Princeton University Press.

Rybicki, J. and Hanski, I. (2013). Species-area relationships and extinctions caused by habitat loss and fragmentation. Ecology Letters, 16:27–38.

Sabo, J. L., Sponseller, R., Dixon, M., Gade, K., Harms, T., Heffernan, J., Jani, A., Katz, G., Soykan, C., Watts, J., et al. (2005). Riparian zones increase regional species richness by harboring different, not more, species. Ecology, 86(1):56–62.

Segatto, P. L., Battin, T. J., and Bertuzzo, E. (2020). Modeling the coupled dynamics of stream metabolism and microbial biomass. Limnology and Oceanography, 65(7):1573–1593.

Simberloff, D. and Gibbons, L. (2004). Now you see them, now you don’t!—population crashes of established introduced species. Biological Invasions, 6:161–172.

Speirs, D. C. and Gurney, W. S. (2001). Population persistence in rivers and estuaries. Ecology, 82(5):1219–1237.

Stott, I., Franco, M., Carslake, D., Townley, S., and Hodgson, D. (2010). Boom or bust? A comparative analysis of transient population dynamics in plants. Journal of Ecology, 98(2):302–311.

Stott, I., Townley, S., and Hodgson, D. J. (2011). A framework for studying transient dynamics of population projection matrix models. Ecology Letters, 14(9):959–970.

Strayer, D. L., D’Antonio, C. M., Essl, F., Fowler, M. S., Geist, J., Hilt, S., Jarić, I., Jöhnk, K., Jones, C. G., Lambin, X., et al. (2017). Boom-bust dynamics in biological invasions: Towards an improved application of the concept. Ecology Letters, 20(10):1337–1350.

Talluto, L., Del Campo, R., Estévez, E., Altermatt, F., Datry, T., and Singer, G. (2024). Towards (better) fluvial meta-ecosystem ecology: A research perspective. npj Biodiversity, 3(1):3.

Terui, A., Ishiyama, N., Urabe, H., Ono, S., Finlay, J. C., and Nakamura, F. (2018). Metapopulation stability in branching river networks. Proceedings of the National Academy of Sciences, 115(26):E5963–E5969.

Tilman, D., May, R. M., Lehman, C. L., and Nowak, M. A. (1994). Habitat destruction and the extinction debt. Nature, 371(6492):65–66.

Townley, S. and Hodgson, D. J. (2008). Erratum et addendum: Transient amplification and attenuation in stage-structured population dynamics. Journal of Applied Ecology, 45(6):1836–1839.

Trevisin, C., Mari, L., Gatto, M., Colizza, V., and Rinaldo, A. (2025). Epidemiological indices with multiple circulating pathogen strains. Infectious Disease Modelling, 10(3):802–812.

Trevisin, C., Mari, L., Gatto, M., and Rinaldo, A. (2024). Epidemicity indices and reproduction numbers from infectious disease data in connected human populations. Infectious Disease Modelling, 9(3):875–891.

Van Looy, K. and Piffady, J. (2017). Metapopulation modelling of riparian tree species persistence in river networks under climate change. Journal of Environmental Management, 202:437–446.

Vannote, R. L., Minshall, G. W., Cummins, K. W., Sedell, J. R., and Cushing, C. E. (1980). The river continuum concept. Canadian Journal of Fisheries and Aquatic Sciences, 37(1):130–137.

Verdy, A. and Caswell, H. (2008). Sensitivity analysis of reactive ecological dynamics. Bulletin of mathematical biology, 70:1634–1659.

Vesipa, R. and Ridolfi, L. (2017). Impact of seasonal forcing on reactive ecological systems. Journal of Theoretical Biology, 419:23–35.

Vincenzi, S., Crivelli, A. J., Jesensek, D., and De Leo, G. A. (2012). Translocation of stream-dwelling salmonids in headwaters: Insights from a 15-year reintroduction experience. Reviews in Fish Biology and Fisheries, 22(2):437–455.

Vincenzi, S., Mangel, M., Jesensek, D., Garza, J. C., and Crivelli, A. J. (2016). Within-and among-population variation in vital rates and population dynamics in a variable environment. Ecological Applications, 26(7):2086–2102.

Vincenzi, S., Mangel, M., Jesensek, D., Garza, J. C., and Crivelli, A. J. (2017). Genetic and life-history consequences of extreme climate events. Proceedings of the Royal Society B: Biological Sciences, 284(1848).

Yeakel, J. D., Moore, J. W., Guimarães Jr, P. R., and de Aguiar, M. A. (2014). Synchronisation and stability in river metapopulation networks. Ecology Letters, 17(3):273–283.

