## Supplementary information for "Reactive persistence of riverine metapopulations"

Lorenzo Mari[a,*], Enrico Bertuzzo[b], Andrea Rinaldo[c,d], Marino Gatto[a], Renato Casagrandi[a]

[a]Dipartimento di Elettronica, Informazione e Bioingegneria, Politecnico di Milano,
Via Ponzio 34/5, Milano 20133, Italy

[b]Dipartimento di Scienze Ambientali, Informatica e Statistica, Università Ca' Foscari Venezia,
Via Torino 155, Venezia Mestre 30172, Italy

[c]Laboratory of Ecohydrology, École Polytechnique Fédérale de Lausanne,
Station 2, Lausanne 1015, Switzerland

[d]Dipartimento di Ingegneria Civile, Edile e Ambientale, Università di Padova,
Via Marzolo 9, Padova 35131, Italy

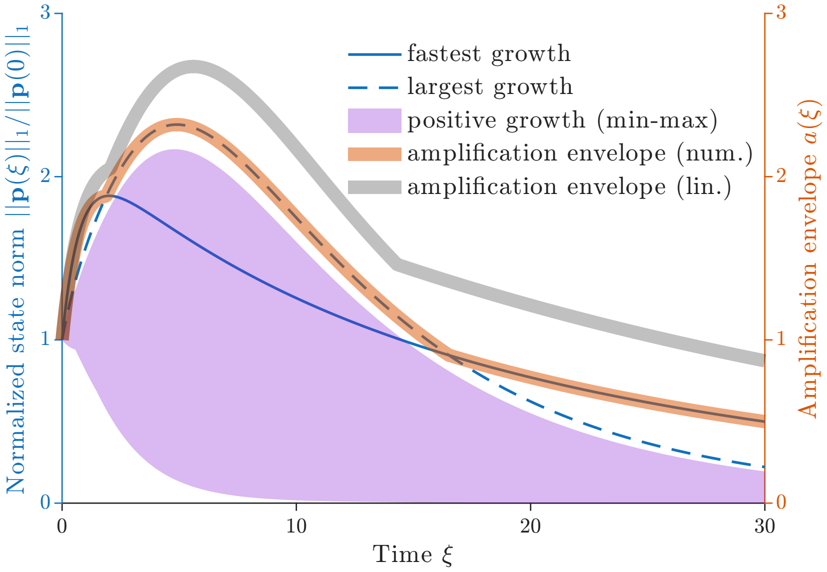


Figure S1: Simulation results for full model (2) in the main text subject to impulsive perturbations to an asymptotically stable, reactive extinction equilibrium. Shown are the temporal dynamics of the normalized $\ell^1$-norm of the state vector (proportional to average patch occupancy), as determined by different initial perturbations (blue lines and purple shading, left axis), and amplification envelope (red thick line, right axis). For numerical simulations, initial conditions are set proportionally to the structure of the fastest-growing perturbation ($\mathbf{p}_\mathbf{0}^\mathbf{f}$, solid blue line), the largest-growing perturbation ($\mathbf{p}_\mathbf{0}^\mathbf{l}$, dashed blue lines), or any other perturbation producing a transient growth in patch occupancy over at least some time scale (purple shading). All configurations have been identified with the linearized form of model (2) in the main text. In all simulations, an initial perturbation of size $||\mathbf{p}(0)|| = 0.1$ has been applied. The amplification envelope has been evaluated numerically considering all single-reach perturbations that lead to a transient growth in average patch occupancy over at least some time scales. The amplification envelope obtained with the linearized counterpart of model (2) in the main text is reported for reference (gray thick line, right axis; see also Fig. 1 in the main text). Parameter values and other details as in Figure 1 in the main text.

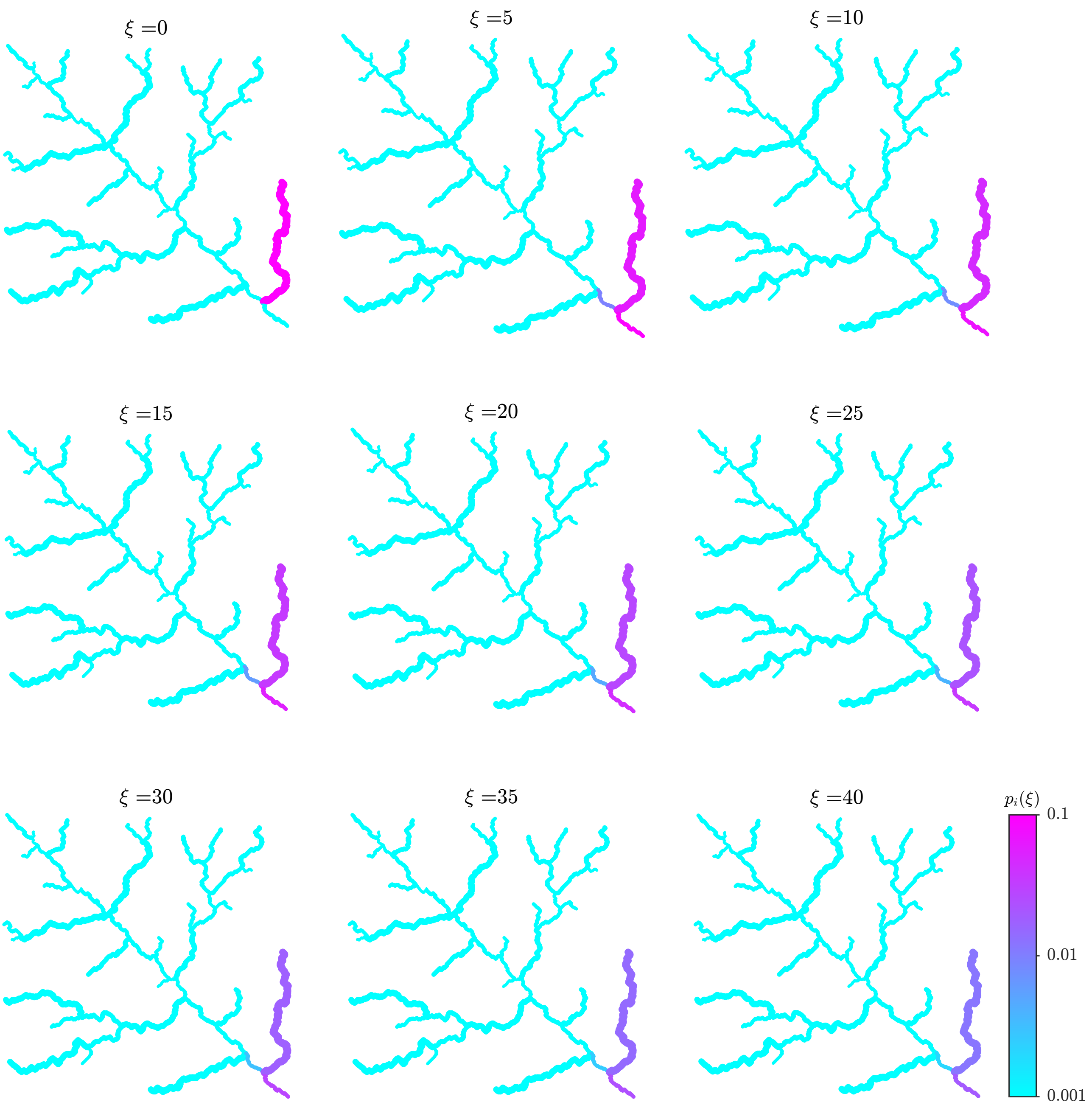


Figure S2: Spatiotemporal patterns of patch occupancy simulated with model (2) in the main text initialized with the fastest-growing perturbation at $\xi = 0$ ($\mathbf{p}_\mathbf{0}^\mathbf{f}$, with $p_j(0) = 0.1$ for $j = j^\mathrm{f}$ and $p_j(0) = 0$ for $j \neq j^\mathrm{f}$). Parameter values and other details as in Figure 1 in the main text.

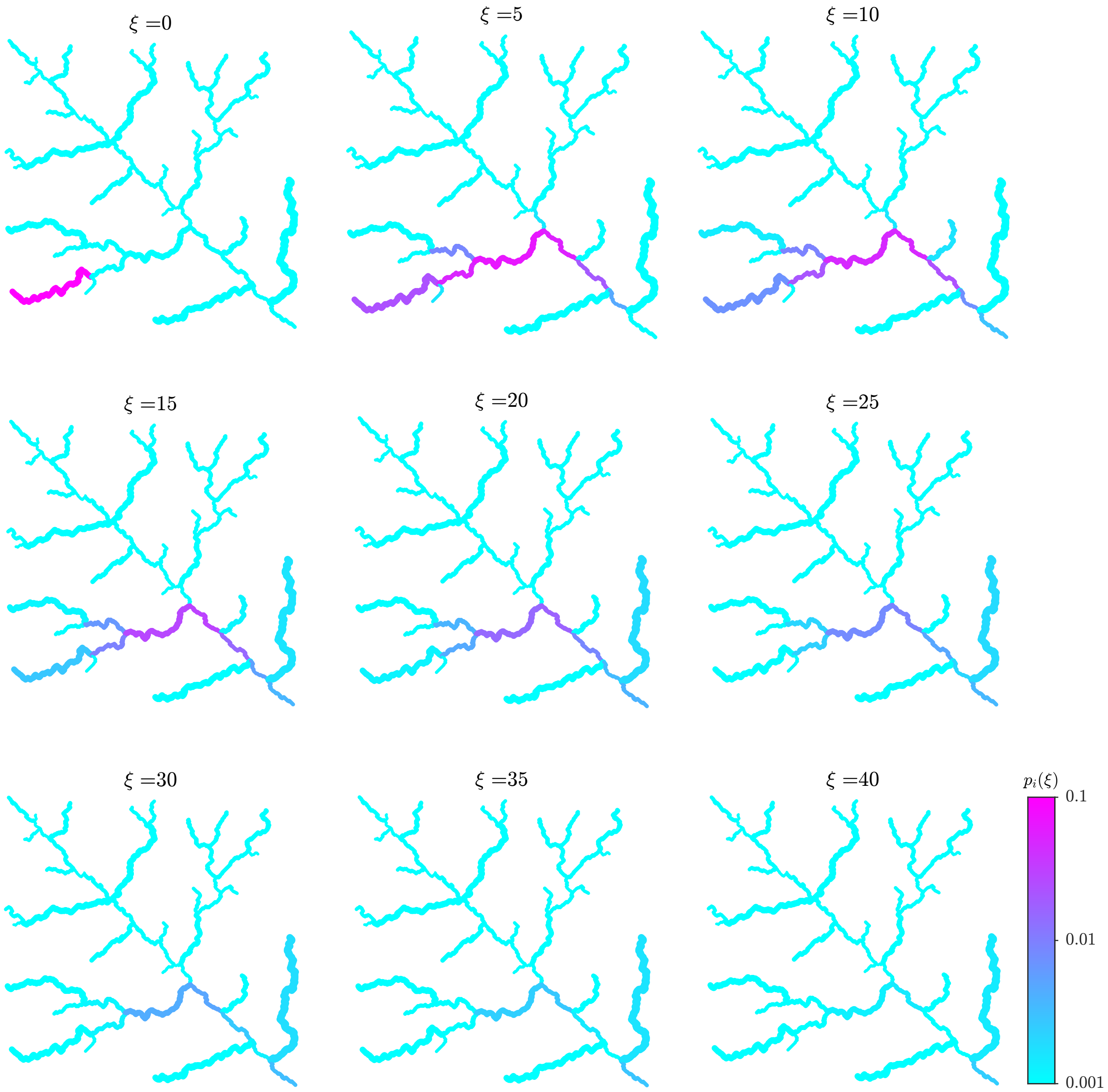


Figure S3: Spatiotemporal patterns of patch occupancy simulated with model (2) in the main text initialized with the perturbation leading to the maximum growth overall ($\mathbf{p_0^l}$, with $p_j(0) = 0.1$ for $j = j^l$ and $p_j(0) = 0$ for $j \neq j^l$). Parameter values and other details as in Figure 1 in the main text.

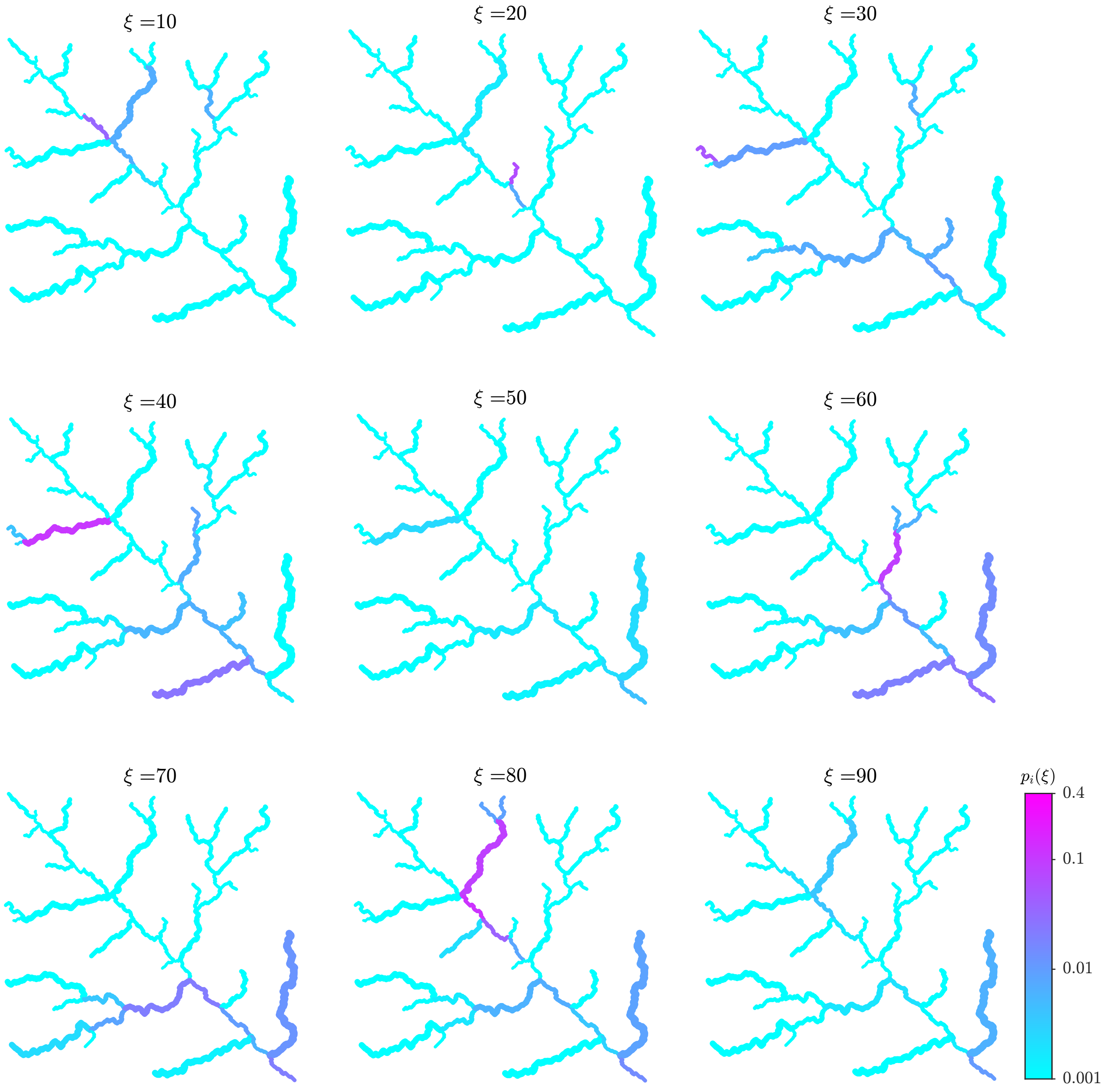


Figure S4: Spatiotemporal patterns of patch occupancy simulated with model (2) in the main text perturbed by recurrent perturbations of the metapopulation extinction equilibrium according to the 'random seeding' scenario considered in Figure 2 in the main text. Only one of the 100 independent realizations of the process is shown. Parameter values as in Figure 1 in the main text, other details as in Figure 2 in the main text.

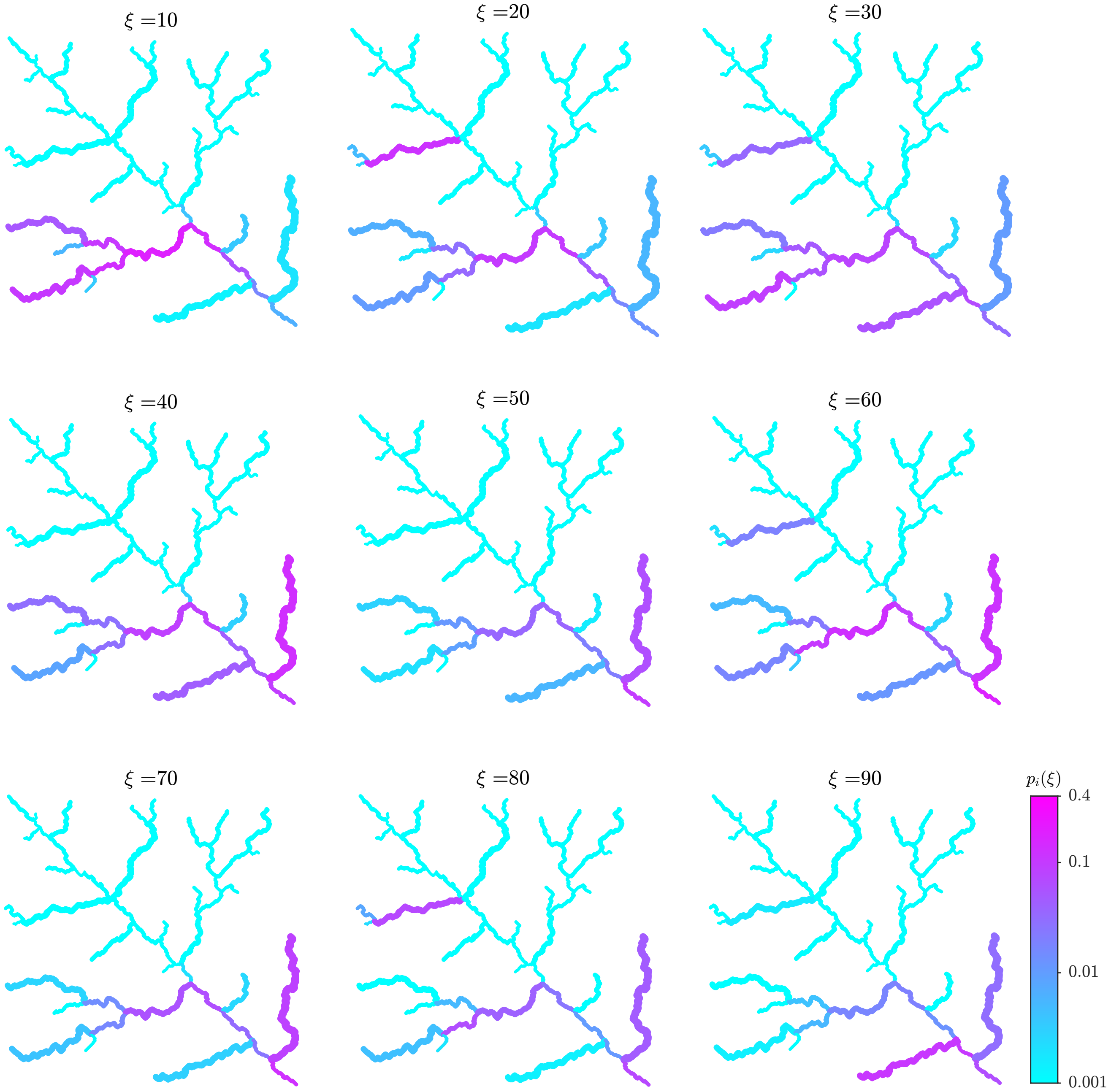


Figure S5: Spatiotemporal patterns of patch occupancy simulated with model (2) in the main text perturbed by recurrent perturbations of the metapopulation extinction equilibrium according to the 'positive growth' scenario considered in Figure 2 in the main text. Only one of the 100 independent realizations of the process is shown. Parameter values as in Figure 1 in the main text, other details as in Figure 2 in the main text.

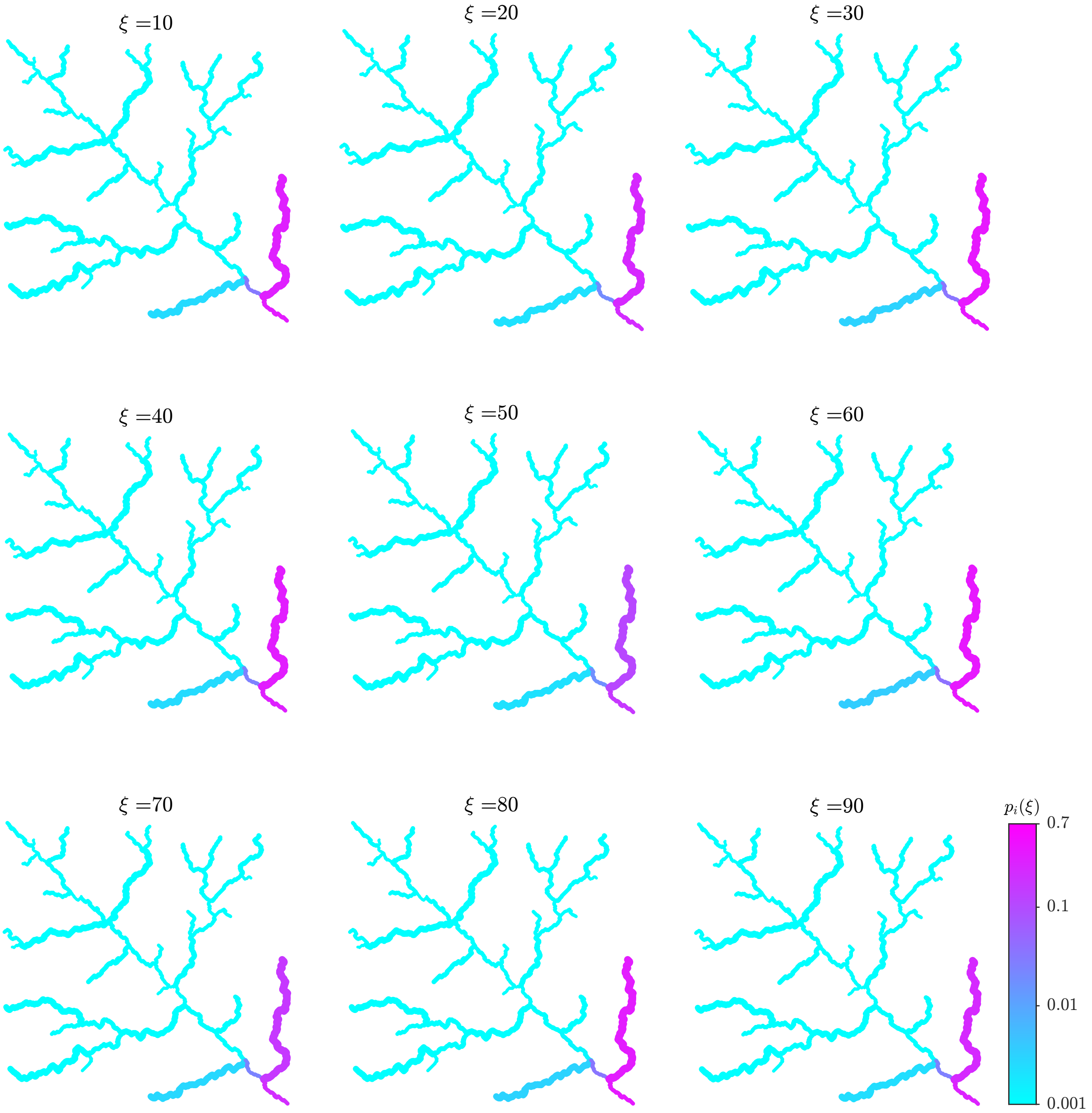


Figure S6: Spatiotemporal patterns of patch occupancy simulated with model (2) in the main text perturbed by recurrent perturbations of the metapopulation extinction equilibrium according to the 'fastest growth' scenario considered in Figure 2 in the main text. Parameter values as in Figure 1 in the main text, other details as in Figure 2 in the main text.

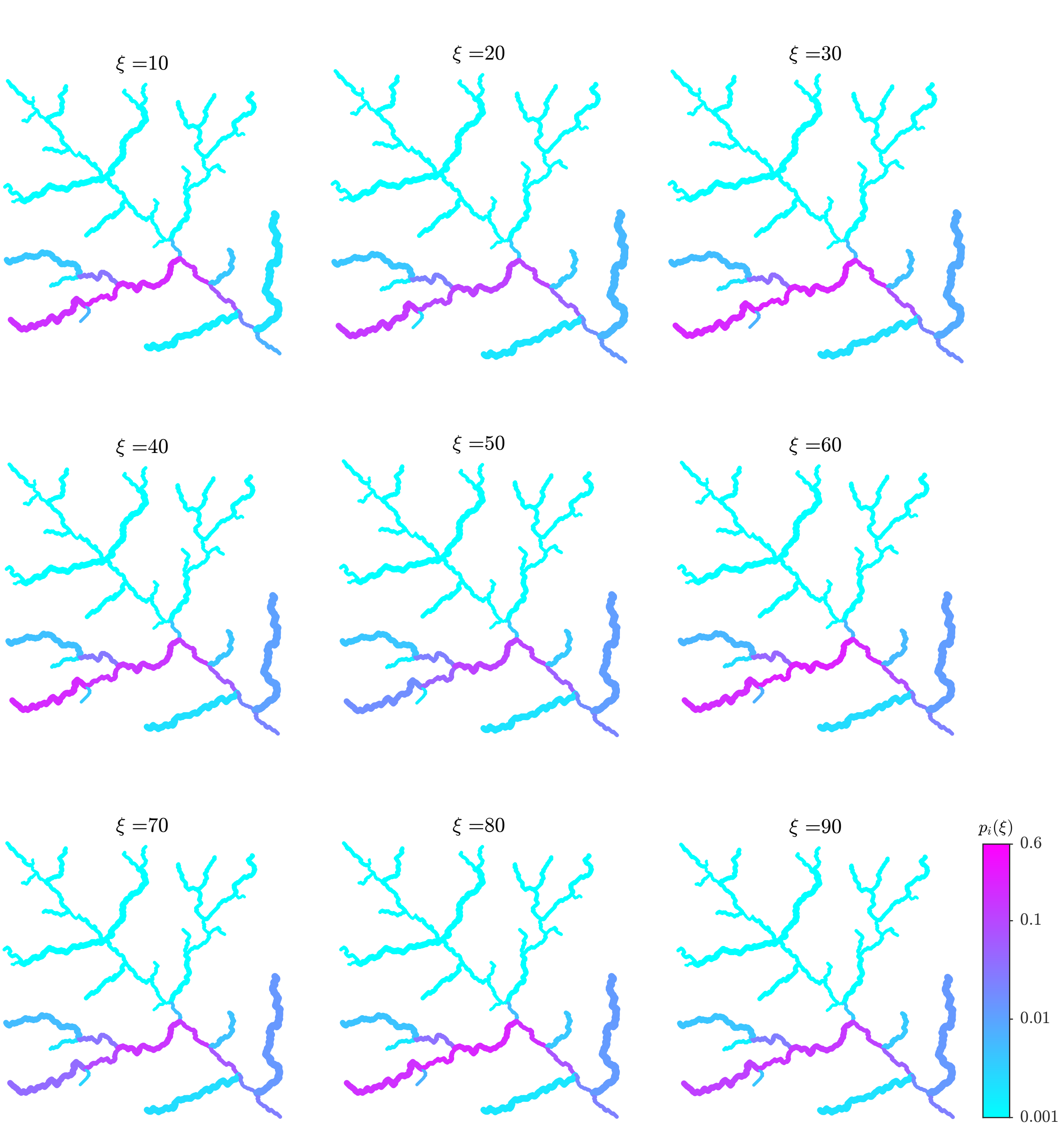


Figure S7: Spatiotemporal patterns of patch occupancy simulated with model (2) in the main text perturbed by recurrent perturbations of the metapopulation extinction equilibrium according to the ‘largest growth’ scenario considered in Figure 2 in the main text. Parameter values as in Figure 1 in the main text, other details as in Figure 2 in the main text.

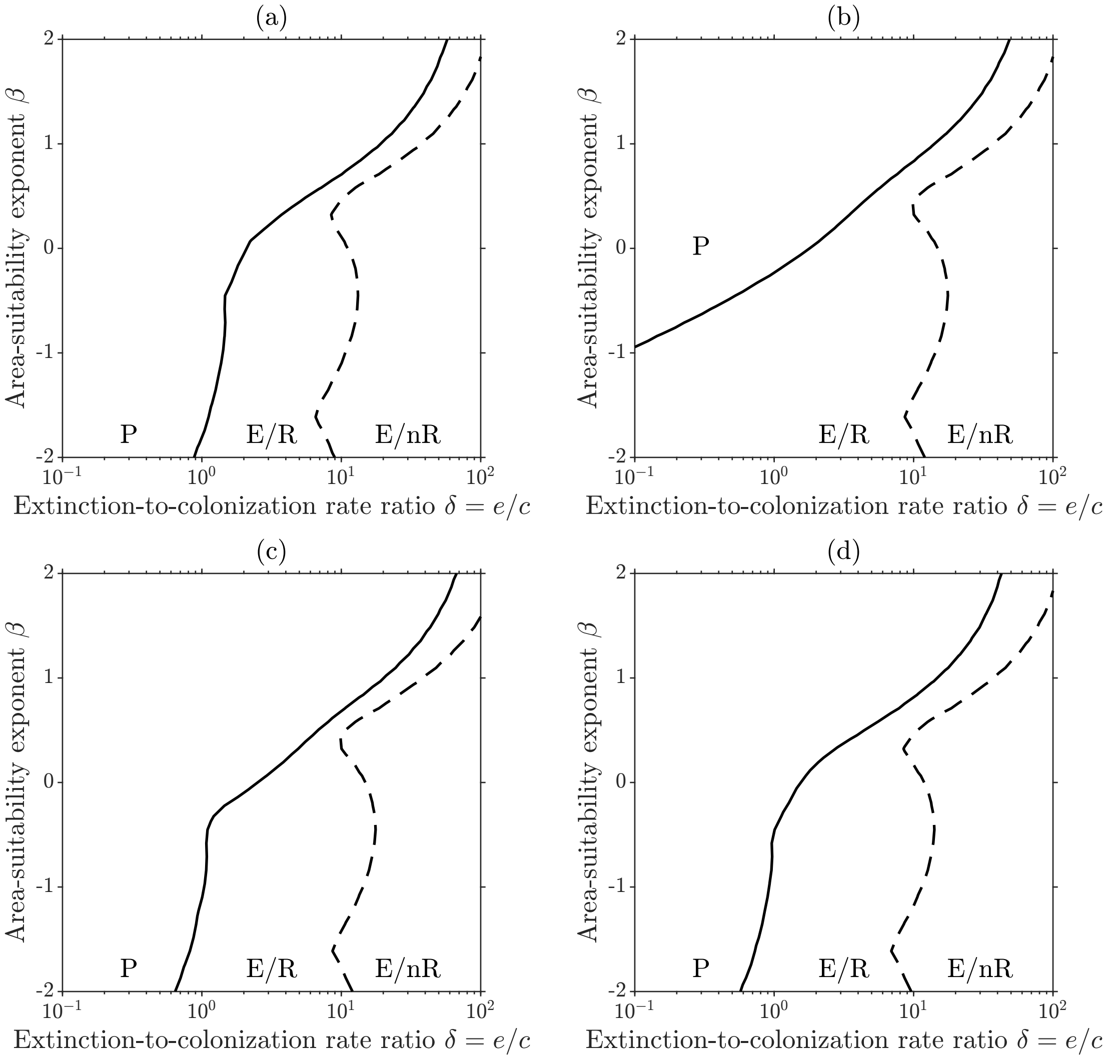


Figure S8: Asymptotic stability and short-term reactivity properties of the extinction equilibrium for alternative parameterizations of dispersal in model (2) in the main text. (a) $\pi = \alpha = 0.5$. (b) $\pi = 1$, $\alpha = 0.5$. (c) $\pi = 0.8$, $\alpha = 0$. (d) $\pi = 0.8$, $\alpha = 1$. In each panel, the solid curve identifies the parameter combinations for which $g = 0$ (or $\rho(\mathbf{M} = \delta)$), which separates the regions (marked as E) in which the extinction equilibrium is asymptotically stable (and the metapopulation cannot persist) from those (marked as P) in which the equilibrium is asymptotically unstable (and the metapopulation can persist). The dashed curve, instead, identifies the condition $r = 0$ (or $\max(\mathrm{CS}(\mathbf{h}) \circ \mathrm{CS}(\mathbf{h}^{-1}\mathbf{M})) = \delta$), which separates the regions (marked as R) where the asymptotically stable extinction equilibrium is reactive (and transient increases in patch occupancy can occur) from those (marked with nR) in which the equilibrium is non-reactive (and transient increases in patch occupancy cannot occur). Other details and parameter values as in Figure 1.

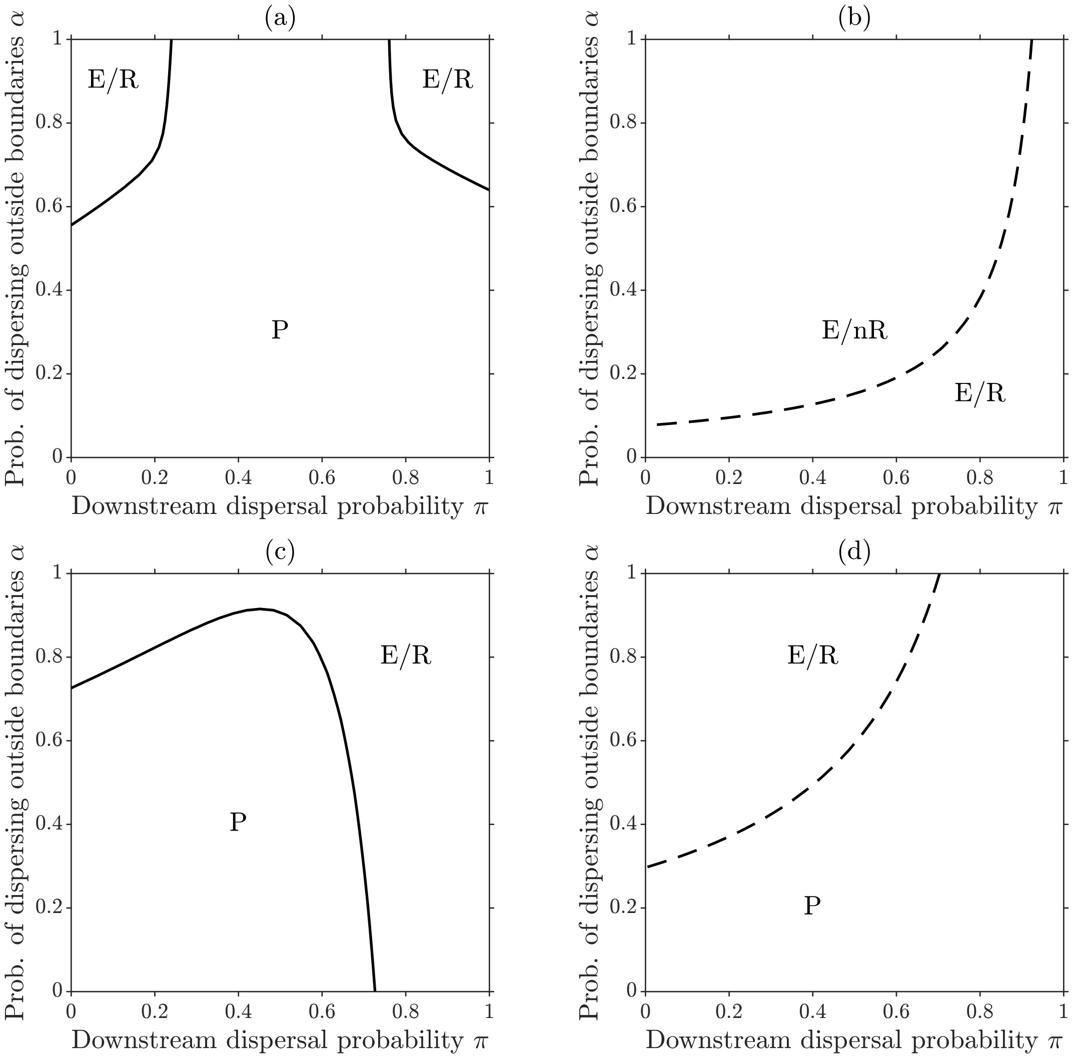


Figure S9: Asymptotic stability and short-term reactivity properties of the extinction equilibrium for alternative baseline parameterizations of the extinction-to-colonization rate ratio and habitat suitability in model (2) in the main text. (a) $\delta = 3$, $\beta = 1/3$. (b) $\delta = 9$, $\beta = 1/3$. (c) $\delta = 1.2$, $\beta = -1$. (d) $\delta = 10$, $\beta = -1$. In each panel, the solid curve identifies the parameter combinations for which $g = 0$ (or $\rho(\mathbf{M} = \delta)$), which separates the regions (marked as E) in which the extinction equilibrium is asymptotically stable (and the metapopulation cannot persist) from those (marked as P) in which the equilibrium is asymptotically unstable (and the metapopulation can persist). The dashed curve, instead, identifies the condition $r = 0$ (or $\max(\mathrm{CS}(\mathbf{h}) \circ \mathrm{CS}(\mathbf{h}^{-1}\mathbf{M})) = \delta$), which separates the regions (marked as R) where the asymptotically stable extinction equilibrium is reactive (and transient increases in patch occupancy can occur) from those (marked with nR) in which the equilibrium is non-reactive (and transient increases in patch occupancy cannot occur). The dots in panel (a) show the parameter combinations explored in panels (b–c), while the dots in panels (b–c) refer to the parameterization of panel (a). Other details and parameter values as in Figure 1.
